# The Endoplasmic Reticulum as a Cytoplasmic Anchor for Microtubule-Driven Aster Centration

**DOI:** 10.64898/2026.10.22.760944

**Authors:** Jérémy Sallé, Dennis Wörthmüller, Javad Najafi, Celia Municio-Diaz, Héliciane Palenzuela, Antoine Rémond, Ilaria Mengoni, Aurélien Guillou, Remi Le Borgne, Catherine Pioche-Durieu, Jean-Marc Verbavatz, Nicolas Joly, Jean François Joanny, Serge Dmitrieff, Nicolas Minc

## Abstract

The proper positioning of microtubule (MT) asters is essential to regulate division positioning and multi-cellular morphogenesis, yet the mechanisms regulating this process remain poorly understood. Using sea urchin zygotes as a model, we combined high-resolution imaging, perturbation assays, and 3D modeling to investigate the nature of forces driving sperm aster centration after fertilization. We show that centering asters are highly anisotropic, featuring long microtubules extending at the aster front that may engage with organelles in bulk cytoplasm to generate pulling forces. While vesicular compartments, like acidic organelles, mitochondria or yolk, contributed minimally, we found that the endoplasmic reticulum (ER) undergoes a massive compaction around the centering aster and drags cytosol to produce a reactive centering force on the aster. Perturbing ER organization by cytoplasmic stratification or Atlastin-mediated fragmentation disrupted aster centration. A two-fluid model suggests that the frictional interaction between the compacting ER and cytosol fluid is sufficient to recapitulate key features of aster centration. These findings identify the ER as an important cytoplasmic anchor for MT force exertion in bulk cytoplasm, to guide aster centering motion and influence early embryonic division geometries. More broadly, our results highlight the role of cytoplasmic architecture in shaping the mechanics of organelle positioning during development.

## INTRODUCTION

The accurate positioning of subcellular structures, like nuclei or mitotic spindles, is fundamental to cell morphogenesis and division. This is particularly important during early embryogenesis, where the position of the nucleus and the orientation of the mitotic spindle determine species-specific cleavage geometries (1). Forces exerted by microtubules (MTs) are well-established regulators of nuclear and spindle positioning (2–5). MTs grow into radial asters from centrosomes that are bound to nuclei or spindles and exert forces that move or position these organelles. Several models for MT force exertion are primarily based on interactions with the cell surface. MTs may for instance generate pushing forces deriving from their polymerization or buckling against the cell surface, which displace nuclei or spindles away from the cortex (6–9). In contrast, cortical pulling MT forces rely on microtubule interactions with molecular motors like dynein anchored at the cell cortex, a mechanism highly relevant to asymmetric divisions to, for instance, decenter the spindle towards a specific cortical polarity domain (10–12). However, a constantly growing body of evidences indicates that interactions between MTs and elements in bulk cytoplasm may also contribute to regulate organelle positioning (13–18). This mode of force exertion may be particularly relevant to the spatial organization and division of zygotes and early blastomeres, which feature a vast cytoplasm. Accordingly, recent assays have shown that the stereotypical motion of centrosomes, asters and nuclei observed during early embryo development can be recapitulated in cytoplasm extracts which lack cellular boundaries and cortices (19–22). To date, however, the mechanisms by which the MT cytoskeleton generates traction forces by interacting with elements in bulk cytoplasm remain fragmented.

One prominent mechanism for MT force generation in the cytoplasm, is the so-called “centrosome-organelle mutual pulling model” (23). This model postulates that the inward-directed motion of vesicular organelles, like endosomes, lysosomes or yolk granules, driven by dynein motors along MTs radiating from centrosomes, yields viscous drag forces that pull MTs and thus centrosomes outward. Given the typical viscosity of the cytoplasm of ∼1 Pa.s and organelle speeds of ∼1µm/s, these drag forces have been proposed to reach significant values of up to ∼1-10 pN (23–25). As longer MTs may engage with more motors or organelles, this model predicts that MT forces may scale with the length of MTs, providing a robust system to probe geometry and target the cell center (23,26–28). This idea was explored and tested in the context of egg fertilization, during which the sperm aster rapidly translocates through the vast egg cytoplasm to reach the cell center, in systems including nematodes, echinoderms, ascidians, fish and frog zygotes (16,23,26,28–31), but recent data have also supported this model in the context of sister aster separation in anaphase (19,32). In addition, this model received support from various modeling approaches (33–35), and its principle was validated in a minimal *in vitro* reconstitution assay (36). However, recent detailed tracking of organelle motion along asters reconstituted in frog cytoplasm extracts, showed that asters and vesicular networks co-move together, questioning the role of vesicular organelle motion in regulating MT force exertion in bulk cytoplasm (19). Furthermore, from a physical point of view, the question of how the cytosol may be advected during organelle/aster movement, and thus how much of the organelle drag force may actually be transmitted to MTs, have raised important limits to this model (15,33,37). Finally, several lines of experiments *in vivo* and *in vitro*, have supported a mechanism more based on MT pushing forces for sperm aster centration in large cells (8,38,39). Therefore, there is an important need to re-explore the mechanism by which MTs could pull from bulk cytoplasm and address its relevance to aster motion in early embryos.

To fill this gap, we revisited the fertilization of sea urchin eggs to address in more detail how astral MTs organize to occupy cytoplasmic space and interact with specific compartments to drive sperm aster centering motion. Using a combination of fixed and live high-resolution imaging, we show that centering asters exhibit a non-homogeneous distribution of MTs, including very long MTs at the aster front that could support cytoplasmic pulling. Moreover, we identify the ER as a critical cytoplasmic anchor for aster motion, as it undergoes a massive inward accumulation during aster centration. This is supported by the observation that redistributing or fragmenting the ER systematically disrupts aster centering trajectories. In addition, we recapitulated aster centration and ER centripetal accumulation using two-fluid finite element simulations, that model aster centering forces as a result of the frictional interaction between the accumulating ER and cytosol. Taken together, our results suggest that aster centering motion requires an intact ER network, which serves as a cytoplasmic anchor for MT force exertion in bulk cytoplasm. These findings highlight how the composition and organization of the cytoplasm may influence sub-cellular positioning forces, thereby shaping division geometries and early embryonic development.

## RESULTS

### The centering aster is highly anisotropic

Sperm aster centration is a remarkably fast and reproducible process, completed within ∼15 minutes (8,26). Shortly after fertilization, microtubules (MTs) start to polymerize from the centrosome brought in the egg by the sperm, to form a sperm aster. In sea urchin embryos, as in many other embryos, the male pronucleus remains close to the centrosome throughout centration, while the randomly positioned female pronucleus is captured upon contacting astral MTs and is transported to the aster center, where karyogamy occurs (39,40). Once centered, the position of the aster center defines the first cleavage plane which determines the symmetric nature of the first embryonic division.

One fundamental ingredient needed to support a model in which the sperm aster is propelled by MT traction forces in bulk cytoplasm stands in a detailed analysis of MT distribution around the aster. Indeed, for this mode of force exertion to account for sperm aster centration, the aster needs to feature an anisotropic shape with longer MTs at its front that can pull more than shorter ones at the back. In addition, these MTs must be far enough from the cortex to rule out putative contributions from cortical pulling. However, capturing the fine details of sperm aster morphology has long been a technical challenge, mainly due to optical limitations in resolving individual MTs and the difficulty of achieving a homogeneous labeling within large egg volumes. We therefore optimized MT visualization methods: For fixed samples, this was achieved by removing the vitelline layer before fertilization, extending permeabilization, and employing secondary nanobodies to enhance immunolabeling quality (see Materials and Methods). This new staining protocol was combined with Airyscan microscopy to permit single MT imaging. Using this approach, we obtained a high signal-to-noise ratio which allowed to track individual MTs within centering asters, often from the centrosome up to the plus ends (**Fig 1A**). Consistent with prior observations, at 5 min post fertilization the aster formed a non-homogeneous radial array of MTs, with the highest density at the aster back, close to the sperm entry point, and a minimum density at the aster front (**Fig 1A-D**) (8,26). Importantly, this enhanced imaging enabled a clearer visualization of full-length MTs in the central cytoplasmic region, where signal was previously diffuse in former staining protocols (8,26). For instance, long MTs or MT bundles up to 52 µm (maximum length = 34.6 ± 8.6 µm, *mean ± SD*) were detected at the aster front whereas those at the rear were significantly shorter (maximum length = 20.7 ± 2.9 µm, *mean ± SD*) (**Fig 1A, E**).

**Figure 1:**
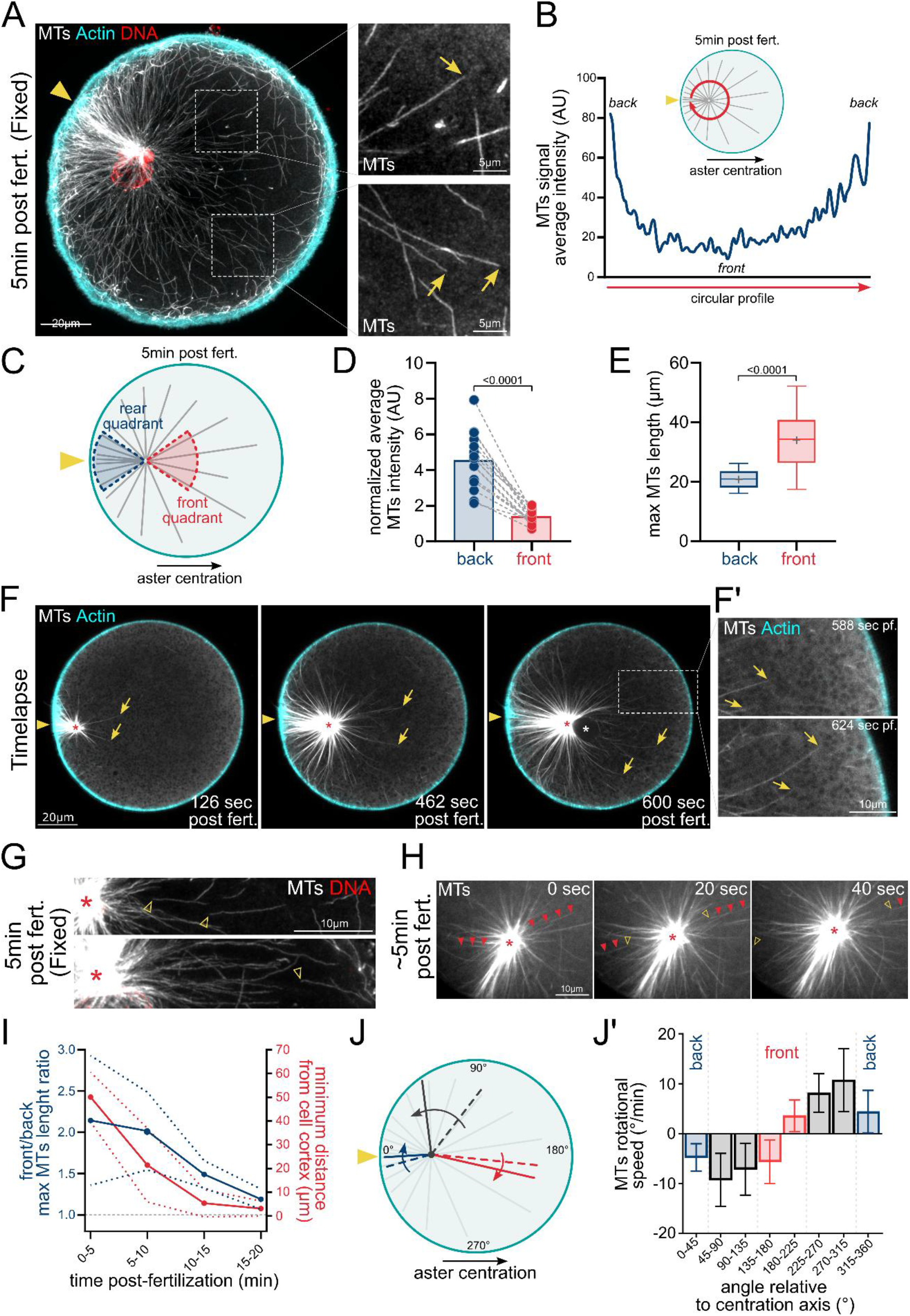
Characterization of the centering aster morphology. (A) Maximum intensity z-projection (10 µm) of confocal images showing immunostaining of a sea urchin zygote 5 min after fertilization. Microtubules are shown in white, actin in cyan, and DNA in red. The yellow arrowhead marks the fertilization site. Scale bar, 20 µm. Right panels show magnified views of two cytoplasmic regions at the aster front; yellow arrows indicate microtubule plus ends. Scale bar, 5 µm. (B) Circular intensity profile of microtubule staining from the image in A. (C) Diagram illustrating the definition of front and rear quadrants (45° angles). (D) Paired quantification of normalized microtubule signal intensity in the rear and front quadrants. Statistical comparison was performed using a paired two-tailed Mann–Whitney test (n = 17 zygotes). (E) Quantification of the average length of the 4–5 longest microtubules in the rear and front quadrants. Statistical comparison was performed using an unpaired two-tailed Mann–Whitney test (n = 156 microtubules from 17 zygotes). (F) Time-lapse images of a sea urchin zygote injected with TMBD-mStayGold and UtrCH-CF555 proteins to label microtubules (white) and actin (cyan), respectively. The yellow arrowhead marks the fertilization site. Yellow arrows indicate microtubule plus ends; the red star marks the aster center, and the white star marks the female pronucleus. Scale bar, 20 µm. (F’) Magnified views of two time points from a cytoplasmic region at the aster front. Scale bar, 10 µm. (G) Examples of immunostainings 5 min after fertilization showing microtubules detached from the aster (yellow arrowheads). Microtubules are shown in white, DNA in red, and the red star marks the aster center. Scale bar, 10 µm. (H) Time-lapse images of microtubules during aster centration (TMBD-mStayGold injection) showing individual microtubules detaching from the aster (arrowheads). Yellow arrowheads mark the minus ends, and red arrowheads mark the rest of the detached microtubules. Scale bar, 10 µm. (I) Evolution of the front/rear ratio of maximum microtubule length during aster centration (left axis), together with the minimum distance between microtubule tips at the aster front and the cell cortex (right axis). The solid line represents the mean, and dotted lines indicate the standard error (n = 5 zygotes). (J) Analysis of microtubule rotational speed around the aster center. The diagram illustrates how radial microtubule orientation changes during aster centration. (J’) Bar plot showing average microtubule rotational speed relative to their initial angle with respect to the centration axis. Data correspond to 142 individual microtubules and were collected from 14 zygotes.

To confirm these observations in live samples, we designed a new MT live probe, achieved by combining the Tau microtubule-binding domain (TMBD) with the mStayGold fluorescent protein (8,41,42). Live imaging of eggs injected with this probe, confirmed the presence of long MTs polymerizing at the aster front, with a ratio of maximum MT length at the aster front versus rear greater than 2 during early centration, similar to that quantified in fixed samples (**Fig 1F**, **Movie1**). Importantly, this ratio gradually decreased to eventually reach a value close to 1 once aster centration was completed (**Fig 1I**). In addition, the minimum distance between long MTs plus ends at the aster front and the cell cortex was ∼50 µm on average during early centration, ∼20 µm between 5-10 min post-fertilization when the aster translocates at constant speed, and only reached values close to 0 µm at 15 min, when the aster finished centration, ruling out putative contributions from cortical MT pulling at the aster front (**Fig 1I**) (8,26). Finally, both live imaging of single MTs and fixed samples, revealed several instances of MTs detaching from the centrosome and gliding away from the aster center, indicating they may be subjected to outward pulling forces in bulk cytoplasm (**Fig 1G-H**, **Movie2**). Together these results appear compatible with a model for aster centration based on MT pulling forces originating from bulk cytoplasm.

The presence of a higher density of MTs at the aster rear, in the form of a “tail” previously served to support a MT pushing mechanism driving aster centration (8). We thus investigated the plausible mechanisms driving tail formation. We first injected an EB1-GFP construct to label growing MT plus ends. Tracking of these plus ends, revealed no major asymmetry in the number of MTs growing from both sides of asters, or in their polymerization rates, suggesting that tail formation may not result from a preferential growth of MTs towards the aster rear (**Fig S1**, **Movie3**). Rather, live imaging of MTs labeled with TMBD-mStayGold, indicated that many MTs tended to pivot backward around their anchoring point at the centrosome, to gradually accumulate at the aster rear (**Fig 1F**, **Movie1**). Accordingly, MT rotational speeds were the highest on the lateral sides of the aster, and always directed towards the aster rear (**Fig 1J**). Such rotational behavior could in part result from MTs being anchored at lateral cortices that rotate as a result of aster forward motion, or from MT frictional interactions with bulk cytoplasm that favor MT alignment along the aster centering path (43). Although these analyses cannot preclude the existence of sperm entry derived elements that may also promote MT stability or bundling at the aster rear, they suggest that aster tail formation may rather be a consequence than a cause of aster centering motion.

### Alternative forces contributing to aster motion

Aster centration could be the outcome of a complex interplay between several types of positioning forces. Among these, as introduced above, cortical pushing forces generated by MT polymerization at the rear may serve as an alternative to cytoplasmic pulling, though these two mechanisms may not be mutually exclusive. Accordingly, in another sea urchin species (*Lytechinus pictus*), experiments using chicken p150^CC1^ protein fragment injection to inhibit dynein activity (44) and suppress MT cytoplasmic pulling showed that asters could still reach the cell center (8). Using the same p150^CC1^ construct at a comparable concentration (≈0.25 µg/ml), we observed that, although zygotes exhibited failed pronuclear fusion and partial cytokinetic furrow ingression, aster centration was indeed not significantly affected (**Fig S2A**). However, capture of the female pronucleus, that serves as a direct readout of dynein-mediated inward transport along MTs (45), occurred normally, indicating that dynein activity may not be properly suppressed during aster centration at this concentration. Therefore, in order to achieve a more rapid and efficient dynein inhibition, we substantially increased the concentration of injected p150^CC1^ (≈50-100 µg/ml). At this concentration, we consistently observed abnormal aster centering behavior, including frequent failures of pronuclear fusion and global aster rotation at late time points (>15 min post-fertilization) (**Movie4**). More importantly, aster trajectories fell into two categories: in 45.5% of cases (n = 11) they were interrupted before reaching the cell center (failed centration) and in the remaining 54.5% of cases they extended into the opposite half of the cell (overshoot) (**Fig S2B-C and S3E**, **Movie4**). Female pronucleus capture also displayed variable behaviors, ranging from wild-type-like motion to delayed or aborted capture. We interpreted these results as evidence of partial dynein inhibition and hypothesized that the limited effect might stem from the use of a non-echinoderm p150^CC1^ sequence. To test this, we generated another p150^CC1^construct based on the *P. lividus* sequence (*Pliv*-p150^CC1^). However, this construct led to essentially similar defects in aster centration and female nuclear capture (**Fig S2C-E**, **Movie4**). Together, these results suggest that the efficacy of p150^CC1^ in inhibiting dynein maybe limited during the short ∼10-15min window of aster centration. Nevertheless, the altered centration phenotypes observed at high p150^CC1^ concentrations support an important contribution of dynein-mediated cytoplasmic pulling forces for aster centration.

Actin polymerization and actomyosin contractility are other well-documented force-generating mechanisms for centrosome positioning, that may play an important role during fertilization and aster motion across species (19,22,46–48). In particular, the acrosome reaction requires the rapid assembly of a sperm-derived actin pool to ensure sperm entry and facilitate early sperm guidance (49). To delineate the contribution of F-actin to MT aster motion, we analyzed early aster centration under various drug treatments. As previously reported, MT depolymerization with nocodazole abolished centration (26). However, the male pronucleus was still able to move inward by an average distance of 5-10 µm (**Fig S3A-B**, **Movie5**), revealing the existence of an initial phase of centrosome motion independent of MTs. Disassembly of F-actin meshworks with latrunculin B resulted in overall normal centration and female nuclear capture, although we noted a slight but reproducible initial lag (**Fig S3A-B**, **Movie5**). When both nocodazole and latrunculin B were applied simultaneously, the male pronucleus remained arrested just beneath the plasma membrane (**Fig S3A-B**, **Movie5**). These results suggest that F-actin contributes modestly during the initial phase of centration, and that loss of actin-mediated forces is largely compensated by MT-dependent forces. Live imaging of MTs and F-actin at the fertilization site supported this interpretation. As previously described (50), we observed local inward growth of F-actin filaments tightly associated with the rear of the centering aster. At approximately 8 minutes post-fertilization, this F-actin structure disassembled as the aster progressed toward the cell center (**Fig S3C and S3E**, **Movie6**). Local F-actin polymerization at the fertilization site persisted after nocodazole treatment, but at a reduced amplitude, indicating that MTs contribute to promote F-actin growth and/or stability during this period (**Fig S3D**, **Movie7**). Taken together, these findings suggest that MT-associated forces are the primary drivers of aster motion, with cytoplasmic forces likely constituting a dominant component of this activity.

### Centering asters generate an inward cytoplasmic flow

To characterize the frictional interaction between MTs and the cytoplasm we next aimed to map cytoplasmic flows during aster centration (32,51,52). If global polarized transport of cytoplasmic elements underlies aster motion, it is likely to generate a detectable cytoplasmic flow in the opposite direction. To isolate the cytoplasmic flow from the net movement of the aster itself we designed a “stalled aster” assay based on the injection of magnetic beads which are used to exert an external force on the aster (25,53). Just before centration was completed, the aster was pulled back toward the cell cortex using magnetic tweezers and held fixed at an off-center position for up to 10 minutes (**Fig 2A**). The flow maps generated using Particle Image Velocimetry (PIV) from the DIC channel revealed a global cytoplasmic flow directed toward the aster center in the central region of the zygote, flanked by vortices caused by cortical backflows (**Fig 2B**, **Movie8**). Although flow velocities exhibited significant variations, the average velocity of the central cytoplasmic flow in the “stalled aster” was approximately 0.005-0.01 µm/s (**Fig 2C**). These results demonstrate the existence of an active inward advection of cytosol along MTs consistent with a cytoplasmic pulling model.

**Figure 2:**
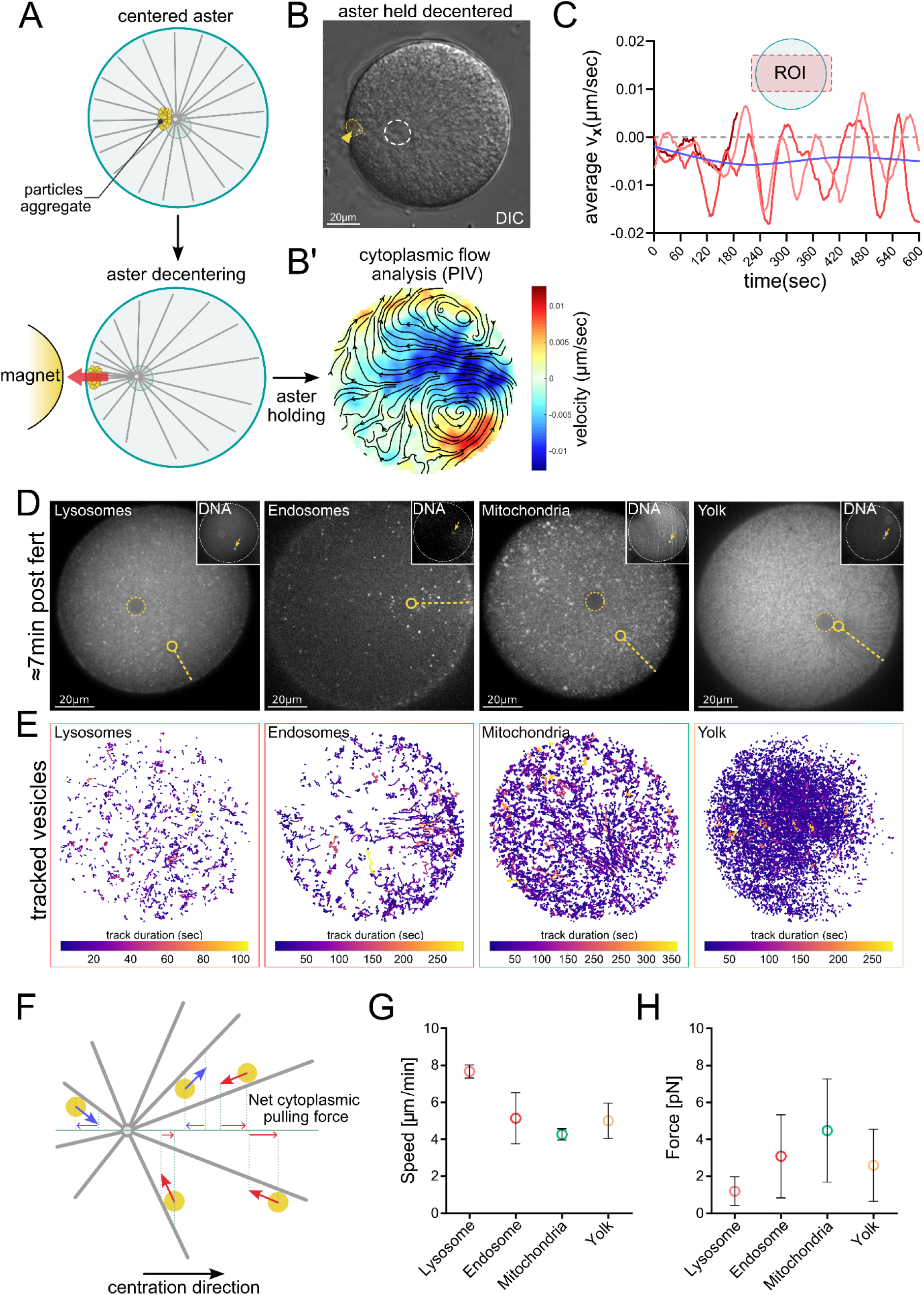
Cytoplasmic flows and spherical vesicle transport during aster centration. (A) Schematic of the “stalled aster” assay in which injected magnetic particles are used to pull and maintain the aster in an off-center position. (B) Representative DIC image of a zygote during the aster-holding phase. The yellow dotted line and arrowhead indicate the magnetic particle aggregate, and the white dotted line marks the position of the nucleus. Scale bar, 20 µm. (B’) Cytoplasmic flow map generated using particle image velocimetry (PIV). The jet colormap represents flow velocity along the centration axis. (C) Average flow velocity over time within the central region of the zygote (n = 3 zygotes). (D) Single frames extracted from time-lapse acquisitions showing, from left to right, lysosomes, endosomes, mitochondria, and yolk granules at approximately 7 min post-fertilization. Yellow dotted lines represent aster trajectories, and yellow circles at their extremities indicate the current position of the aster center. Yellow dotted circles highlight the female pronuclei, when visible in the plane of imaging. Top-left insets correspond to the DNA channel (Hoechst). Yellow arrows indicate the position of the male pronucleus. Scale bar, 20 µm. (E) Corresponding individual vesicles tracks. The color code corresponds to the track duration. For clarity, only trajectories with more than 10 track points are displayed. (F) Scheme illustrating how the net cytoplasmic pulling force was estimated from individual vesicle trajectories. (G) Measured average velocity for each vesicle category. (H) Estimated average net cytoplasmic pulling force for each vesicle category. The dataset includes 15862, 5920, 24409, and 80404 individual tracks for lysosomes, endosomes, mitochondria, and yolk granules, respectively, collected from three independent zygotes per category. Error bars represent SD.

### Friction forces produced by cytoplasmic vesicular compartments are insufficient to promote aster motion

The ooplasm forms a dense, homogeneous mixture of micron-sized spherical organelles, including mitochondria, yolk granules, endosomes, and lysosomes, interwoven with the endoplasmic reticulum (ER) membrane network. Collective minus-end directed motions of various organelles have already been reported in other systems and, in some cases, have been proposed to be one driving mechanisms behind aster centration (23). To address whether aster motion is associated with the inward transport of a specific compartment in our system, we first focused on spherical vesicles, and tracked their trajectories in a single focal plane during aster centration using specific dyes (**Fig 2D-E**, **Movie9**). Interestingly, only a limited fraction of vesicles exhibited fast radial motions relative to aster center that would be compatible with transport along MTs. Most vesicles instead appeared to follow global aster motion, with only weak and transient accumulation around the aster center for endosomes, no accumulation for lysosomes and mitochondria, and even exclusion from the aster center upon completion of centration in the case of yolk granules, in line with findings in frog extracts (**Fig 2D-E**, **Movie9**) (19). To roughly estimate the maximal friction forces generated by the displacement of these spherical compartments, we projected the trajectory of each vesicle onto the aster centration axis (**Fig 2F**). Using measured vesicle velocities, sizes and cytoplasm viscosity, we applied Stokes’ law to estimate the net pulling force exerted on the aster (**Fig 2G-H**). The estimated net force directed toward the cell center was on average less than ≈12 pN, far below previously measured aster stall force of ≈600 pN obtained with *in vivo* magnetic tweezers (25). Overall, these results suggest that the transport of lysosomes, endosomes, mitochondria or yolk granules may not contribute significantly to aster motion in this system.

### A global ER network reorganization is associated with aster centration

The endoplasmic reticulum is classically described as a complex endomembrane network of interconnected tubes and sheets (54). In the sea urchin ooplasm, it represents the second most abundant membrane compartment after yolk granules (55). Because the deformation and displacement of an extended ER network are expected to generate a greater viscous drag force than spherical vesicles, the ER is a solid candidate as a cytoplasmic anchor capable of influencing aster motion (56). To test this possibility, we first characterized ER dynamics during aster centration by labeling ER membranes through the injection of oil droplets saturated by DilDye (57,58). We observed that the ER, initially distributed as a homogeneous cytoplasmic network, rapidly accumulated around the aster center after fertilization, forming a region of increasing density as the aster moved towards the cell center (**Fig 3A-B**, **Movie10**). Following centration, ER accumulation continued, ultimately forming a dense perinuclear domain that remained associated with the center of interphase asters (58). To investigate whether this global ER reorganization was accompanied by changes in network morphology, we performed serial block-face scanning electron microscopy (SBF-SEM) to analyze ER reorganization. Pixel classification-based 3D segmentation of ER, yolk granules and mitochondria revealed that, in the unfertilized egg, ER membranes form a regular mixture of tubes and sheets interspacing yolk granules and mitochondria, all homogeneously distributed throughout the cytoplasm (**Fig 3C-D**). As expected, after aster centration, the ER formed a denser membrane network in the perinuclear region. However, we were not able to clearly detect any obvious change in network morphology (**Fig 3E-F**). In addition, and in agreement with live imaging assays, yolk granules appeared excluded from the vicinity of the aster center (**Fig 3C and 3E**). These observations suggest that the ER undergoes a targeted “compaction” process rather than a wholesale structural reorganization, likely driven by active minus-end–directed transport. To explore this possibility and directly visualize ER transport dynamics, we carried out high–frame-rate imaging following DilDye injection. Remarkably, these revealed short-lived, centripetal ER movements with velocities of 17.3 ± 11.4 µm/s (mean ± SD) and long persistent displacements of 3.6 ± 2.1 µm (mean ± SD), consistent with typical speeds and persistence of dynein motors *in vivo* (59). These rapid, transient motions resulted in localized deformations within the ER membrane network (**Fig 3G-I**, **Movie11**). These discrete pulling events are consistent with the action of MT-associated motors generating inward cytoplasmic pulling forces. Taken together, these results led us to propose a working hypothesis in which aster movement is driven by the cumulative effect of multiple successive pulling events on the ER network by motors associated to MTs in bulk.

**Figure 3:**
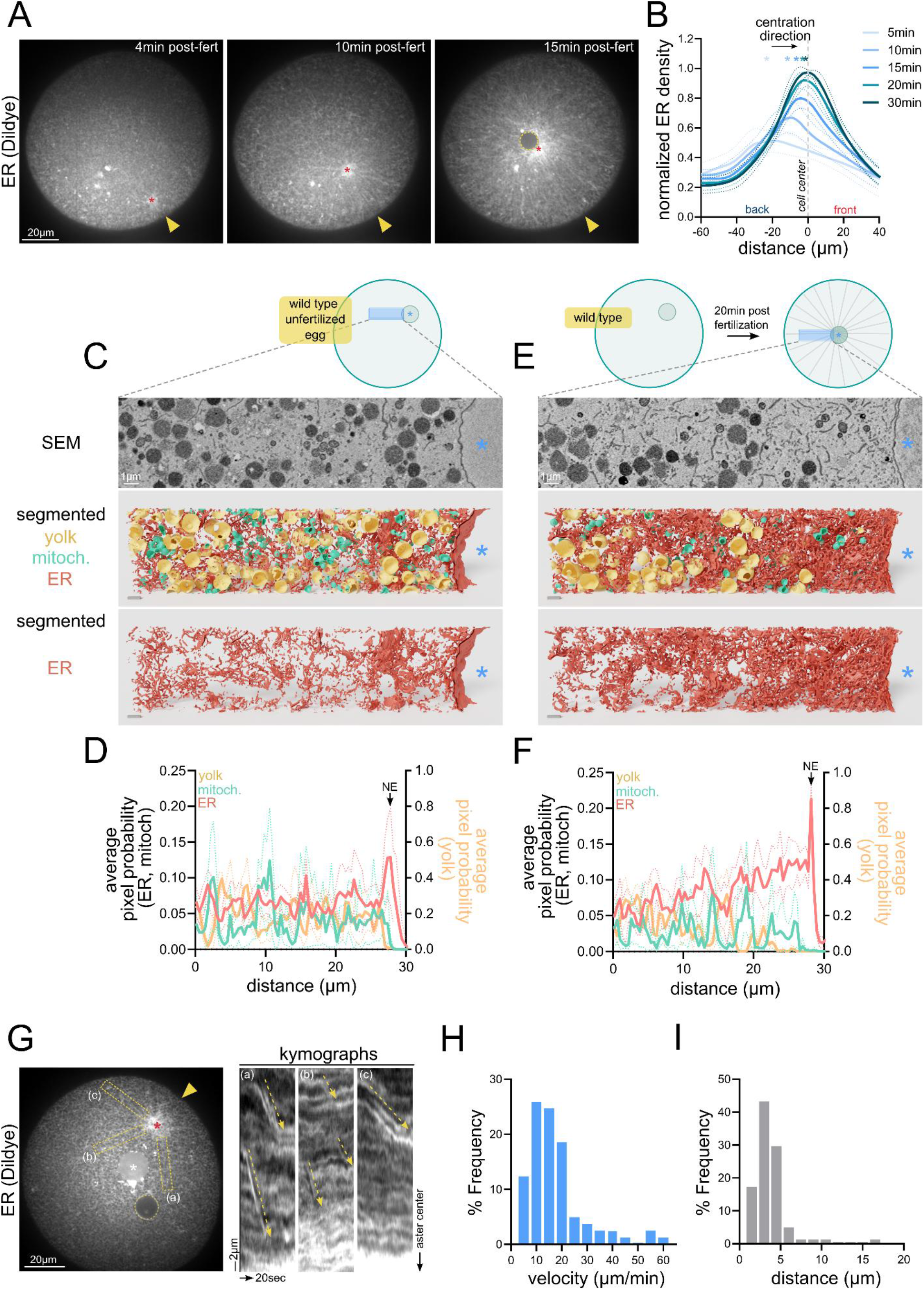
ER network reorganization during aster centration. (A) Time lapse images of aster centration following Dildye injection (ER labelling). The yellow arrowhead indicates the fertilization site, the red star marks the aster center, and the yellow dotted circle outlines the female pronucleus. Scale bar, 20 µm. (B) Normalized ER density dynamics relative to the cell center during aster centration. Each curve corresponds to a specific time point from 5 to 30 min post-fertilization. Dotted lines indicate SD, and overlaid stars mark the position of the aster center relative to the cell center (n = 11 zygotes). (C–E) Cytoplasmic segmentation of a cuboid-shaped volume containing part of the nucleus in an unfertilized egg (C) and at 20 min post-fertilization (E). Raw z-stacks were acquired using serial block-face scanning electron microscopy (SBF-SEM) (top row). Yolk (yellow), mitochondria (green), and ER (red) were segmented using Ilastik software. Pixel probability maps were used to generate 3D renderings of cytoplasmic organization (middle and bottom rows). The blue star marks the nucleoplasm. Scale bar, 1 µm. (D–F) Corresponding analysis of pixel probability profiles for the unfertilized egg (D) and at 20 min post-fertilization (F). Each curve represents the mean pixel probability across all z-sections for the three analyzed compartments. Dotted lines indicate SD. The rightmost peak in the ER profile corresponds to the nuclear envelope (NE). (G) High-frame-rate imaging of ER dynamics following DiIdye injection. The left panel shows a still image from a movie in which three radial cytoplasmic zones (a, b, and c) are highlighted by red dotted boxes. The yellow arrowhead marks the fertilization site, the red star the aster center, the yellow dotted circle the female pronucleus, and the white star the oil droplet (DiIdye). Scale bar, 20 µm. The right panels show kymographs from each cytoplasmic region, with centripetal ER pulling events highlighted by yellow dotted lines. Horizontal and vertical scale bars represent 2 µm and 20 s, respectively. (H–I) Frequency distribution of ER pulling event velocities (H) and distances (I) (n = 81 events from 8 zygotes).

### Cytoplasm stratification biases aster positioning towards ER-enriched domains

To investigate the contribution of the ER to aster positioning, we first aimed to generate cytoplasmic regions with varying densities of ER membranes. Our rationale was that if aster motion was independent of the ER, its positioning should occur similarly in both ER-enriched and ER-depleted regions. Conversely, if the ER contributes to aster movement, we would expect a preferential positioning within ER-enriched domains. Different ER densities were obtained through centrifugation-induced stratification of the cytoplasm (60,61). Stratified eggs adopt a characteristic “hourglass” shape elongated along the centrifugation axis, with the ER mostly restricted to the centripetal half with a higher density near the centripetal pole (**Fig 4A-B**). Other compartments are also redistributed: Like the ER, the female pronucleus localized to the centripetal pole, while mitochondria accumulated at the centrifugal pole, and yolk granules were enriched in the centripetal half but excluded from a small region at the centripetal pole (**Fig 4B**). Following fertilization, stratified zygotes were classified into three groups according to the sperm entry site location (centrifugal, centripetal, or middle zone) (**Fig 4A**). Strikingly, final aster positioning consistently displayed a bias toward the ER-enriched domain, relative to the expected position at the cell geometrical center (26). This bias became more pronounced when fertilization occurred closer to ER-enriched regions, although aster velocity remained constant across all conditions (**Fig 4C-E**). This suggests that ER density influences aster positioning primarily by guiding its trajectory toward the ER-rich domain. As a consequence of aster mis-centering, at the first cleavage, blastomeres displayed a pronounced size asymmetry, reflecting the preceding aster displacement (**Fig 4F**). However, the location of the fertilization site did not seem to influence the amplitude of this asymmetry (**Fig 4F-G**). This could be explained by a progressive re-homogenization of the cytoplasm and/or a partial recentering motion during spindle assembly or anaphase. Although contributions from other cytoplasmic components (*e.g.* acidic organelles) could not be fully excluded from these assays, components enriched at the centrifugal pole, such as mitochondria, appeared to be dispensable. Collectively, these results support the notion that the ER network contributes to aster motion.

**Figure 4:**
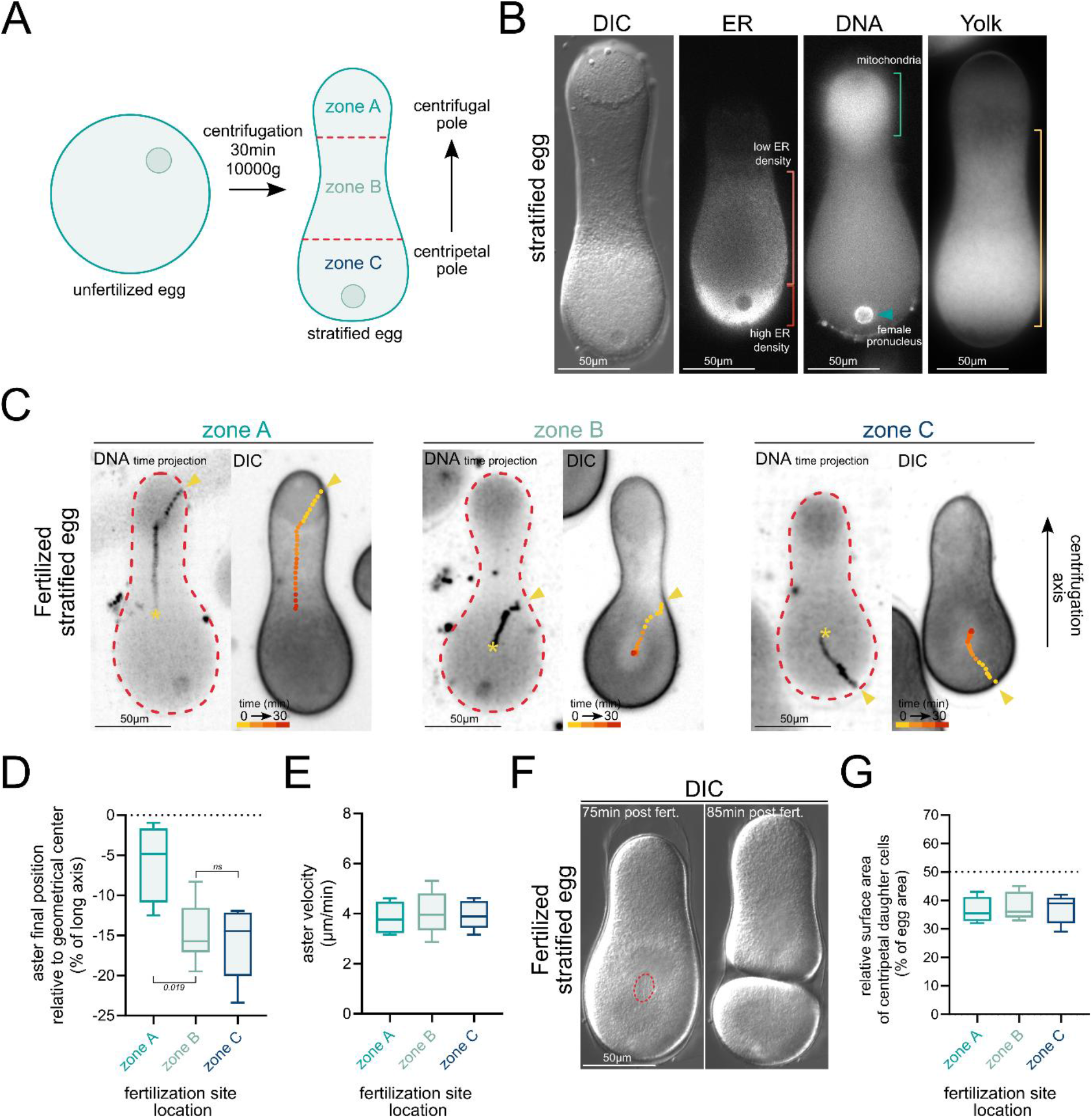
Aster trajectories follow ER-rich regions in stratified eggs. (A) Schematic representation of the cytoplasmic stratification protocol based on centrifugation of unfertilized eggs. Stratified eggs are elongated along the centrifugation axis and can be divided into three regions: a centrifugal region (zone a) enriched in high-density compartments, a central region (zone b), and a centripetal region (zone c) enriched in low-density compartments and containing the female pronucleus. (B) Distribution of ER, mitochondria, and yolk granules in fixed stratified eggs. From left to right: a DIC image of a stratified egg; anti-KDEL immunostaining showing ER enrichment at the centripetal pole (orange and red brackets indicate low and high ER density, respectively); DNA staining (Hoechst) showing the female pronucleus at the centripetal pole and mitochondria accumulation at the centrifugal pole (blue bracket); yolk granule labeling (Nile blue) showing the overall yolk enrichment in the centripetal half of the egg and yolk exclusion from the most centripetal region (yellow bracket). Each image corresponds to a different egg. Scale bar, 50 µm. (C) Representative examples of aster trajectories in stratified eggs fertilized in zone a (left), b (middle), or c (right). For each condition, the left panel shows a time projection of the DNA signal (Hoechst), and the right panel shows a DIC image of the final frame of the time-lapse acquisition with the aster trajectory superimposed (time post-fertilization is color-coded). The yellow arrowhead indicates the fertilization site, and the red star indicates the final aster position. (D) Deviation between the final aster position and the geometric center of the egg, expressed as a percentage of the cell long axis. Statistical comparisons were performed using a two-tailed Mann–Whitney test (p values are indicated on the graph; ns, p > 0.05). (E) Corresponding measurements of aster velocity (n = 4, 9, and 6 for zones a, b, and c, respectively). (F) Representative DIC images from a time-lapse acquisition of the first division of a fertilized stratified egg. Scale bar, 50 µm. (G) Relative surface area of the centripetal daughter cell, expressed as a percentage of the total egg area (n = 4, 11, and 7 for zones a, b, and c, respectively).

### ER fragmentation impairs aster centration

To further examine the contribution of the ER to aster centering, we designed an experiment in which the ER membrane network was altered in a way that would affect its frictional interaction with the cytoplasm. To this end, we used a dominant-negative form of the ER membrane protein Atlastin (ATL) which mediates ER membrane fusion through the homotypic interaction of its GTP-binding cytoplasmic domain (62). This dominant-negative construct consists of the isolated cytoplasmic domain (cytATL) which can bind endogenous Atlastins but lacks the ability to fuse ER membranes, ultimately leading to ER network fragmentation (63). To assess the impact of cytATL in our system we used SBF-SEM to characterize ER network defects. At 20min post-fertilization, embryos injected with a control mutated form of cytATL (cytATL^R232Q^) which is unable to interact with endogenous ATL, displayed the characteristic accumulation profile observed in non-injected control embryos (**Fig 5A and 5C**). In contrast, in cytATL-injected embryos, the central ER accumulation was abolished. Instead, ER density was lowest near the aster center, forming a 30–40 µm exclusion zone, while remaining relatively homogeneous throughout the rest of the cytoplasm (**Fig 5B and 5D**). Notably, the distributions of yolk granules and mitochondria were not significantly affected, suggesting that their exclusion from the aster center may not be driven by ER accumulation (**Fig 5B and 5D**). Closer inspection of ER membranes revealed that the low-density region consisted of numerous small ER vesicles, consistent with the cytATL-induced fragmentation phenotype reported in other systems (**Fig 5B, lower panel**) (63,64).

**Figure 5:**
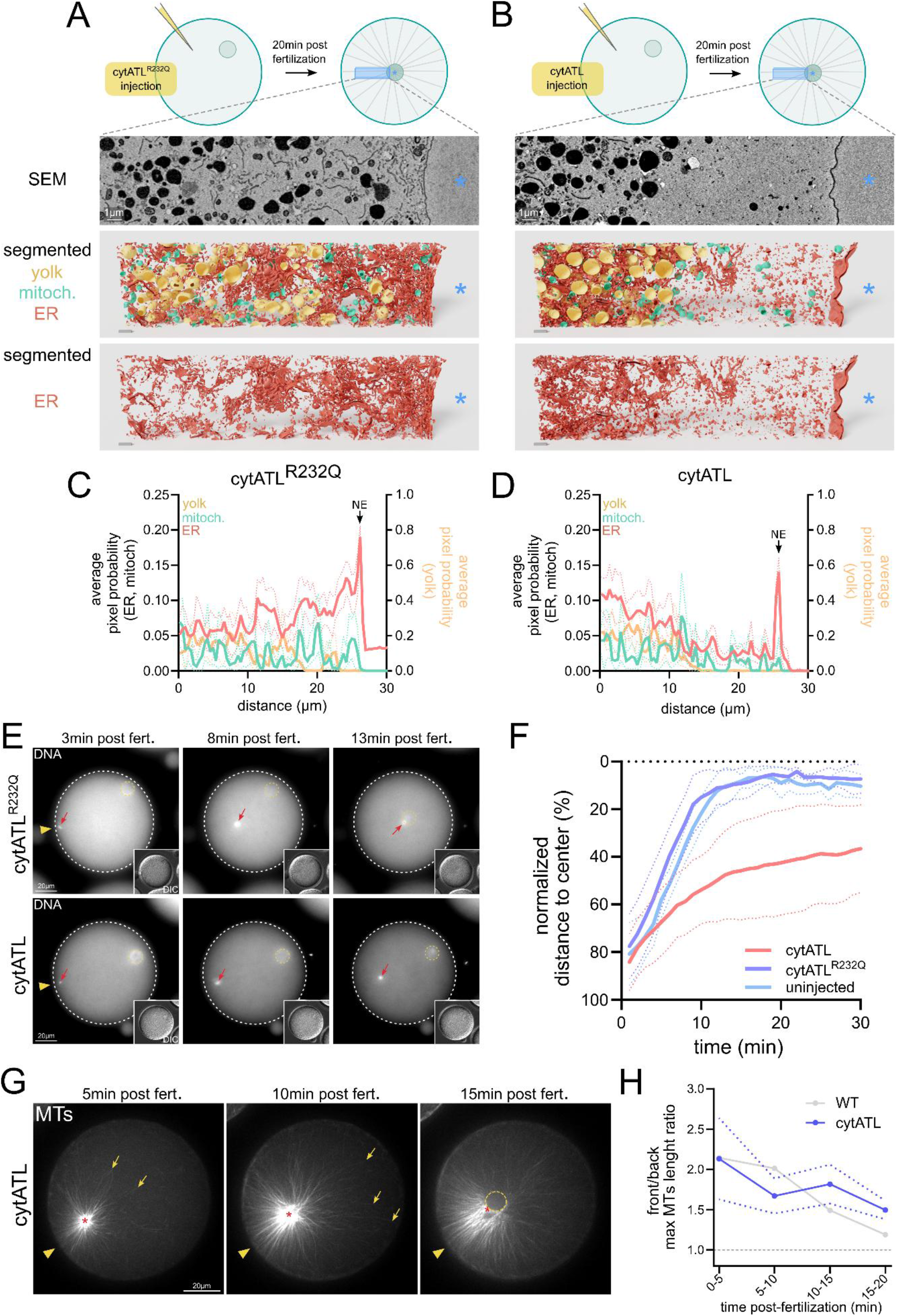
Aster centration requires an intact ER network. (A–B) Cytoplasmic segmentation of a cuboid-shaped volume containing part of the nucleus at 20 min post-fertilization after cytATL^R232Q^ (A) and cytATL protein injection (B). Raw z-stacks were acquired using serial block-face scanning electron microscopy (SBF-SEM) (top row). Yolk (yellow), mitochondria (green), and ER (red) were segmented using Ilastik software. Pixel probability maps were used to generate 3D renderings of cytoplasmic organization (middle and bottom rows). The blue star marks the nucleoplasm. Scale bar, 1 µm. (C–D) Corresponding analysis of pixel probability profiles for the zygote injected with cytATL^R232Q^ (C) and cytATL (D). Each curve represents the mean pixel probability across all z-sections for the three analyzed compartments. Dotted lines indicate the standard error. The rightmost peak in the ER profile corresponds to the nuclear envelope (*ne*). (E) Time-lapse images of aster motion (Hoechst 33342) in zygotes injected with cytATL^R232Q^ (top) and cytATL (bottom) proteins. Insets show the DIC channel. Red arrows show the male pronucleus (aster center) and the yellow dotted circles outline the female pronucleus. Scale bar, 20µm. (F) Dynamics of the normalized average distance between aster center and the cell center in uninjected zygotes (in blue, n= 6), and zygotes injected with cytATL^R232Q^ (in purple, n= 5) or cytATL (in red, n= 18) proteins. Dotted lines indicate the standard error. (G) Time-lapse images of a sea urchin zygote during aster centration injected with TMBD-mStayGold and cytATL proteins to label microtubules (in white) and induce ER fragmentation, respectively. The yellow arrowhead marks the fertilization site. Yellow arrows point at microtubule plus ends; the red star marks the aster center, and the yellow dotted circle marks the female pronucleus. Scale bar, 20 µm. (H) Evolution of the front/rear ratio of maximum microtubule length during aster centration after cytATL protein injection. The solid blue line represents the mean, and the dotted blue lines indicate the standard error (n= 4 zygotes). The grey line corresponds to the mean measured in embryos injected only with TMBD-mStayGold (see Fig1).

Strikingly, injection of purified cytATL protein caused a premature interruption of aster centration, preventing asters from reaching the cell center (**Fig 5E-F**, **Movie12**). Female pronucleus capture was also impaired, with reduced and/or delayed movement, indicating that female pronucleus transport may be associated to ER inward transport. In contrast, cytATL^R232Q^ injections produced aster trajectories comparable to those of non-injected controls (**Fig 5E-F**, **Movie12**). Because microtubule–ER interactions and mutual regulations have been reported (65–67), we sought to confirm that the cytATL phenotype was not a consequence of defective aster growth or morphogenesis. We combined cytATL injection with live MT imaging, and detected no significant deviations from the typical aster architecture, with higher MT density at the rear and longer MTs at the front (**Fig 5G**, **Movie13**). However, since centrosomes remained off-centered beyond 15 minutes after fertilization, the MT front-to-back length ratio stabilized at ∼1.5 instead of converging toward 1, indicating that MT growth can continue even when the ER and centration are perturbed (**Fig 5H**). This observation reinforces the idea that cortical pushing forces alone are insufficient to drive aster centration. Taken together, these results demonstrate that ER disorganization and/or partial fragmentation can halt aster motion, suggesting that the ER membrane network serves as a cytoplasmic anchor that enable MTs to generate traction forces from the cytoplasm.

An interesting feature of the cytATL injection phenotype is the persistence of initial aster movement, whose amplitude is too large to be explained solely by the above-mentioned contribution of F-actin polymerization at the fertilization site. To address this, we analyzed ER network fragmentation dynamics by performing consecutive injections of Dildye and cytATL protein. In unfertilized eggs injected with cytATL, ER organization was initially homogeneous and indistinguishable from that of non-injected eggs. However, after fertilization, an ER-depleted zone containing mostly small ER fragments formed at the aster center and progressively expanded into a ring of ER disconnected from the aster center (**Fig 6A-B**, **Movie14**). Moreover, MT depolymerization with nocodazole in cytATL-injected zygotes prevented ER network fragmentation (**Fig S4A-B, Movie15**), indicating that cytATL-dependent ER fragmentation is caused by MTs within asters which may gradually disrupt the ER network. Interestingly, the formation of an ER-depleted zone allowed us to visualize membrane tubes extending toward the aster center, reminiscent of membrane tubes pulled *in vitro* by MT associated motors like kinesins (68). These tubes, originating from the peripheral ER network or the nuclear envelope, exhibited rapid back-and-forth movements, presumably reflecting transient unproductive pulling events that cause ER tubes to snap back (**Fig 6C, Movie16**). Similar events were observed more frequently at later time-points around spindle poles likely because of the increased density in astral MTs in mitosis (**Fig 6D, Movie17**). Based on these results, we interpret the cytATL phenotype as a progressive fragmentation of a weakened ER network, that cannot repair, caused by failed pulling events, which gradually reduce cytoplasmic pulling efficiency and ultimately prevent aster movement. Therefore, these results reinforce the hypothesis that ER-generated cytoplasmic pulling forces may play a major role in aster centering motion.

**Figure 6:**
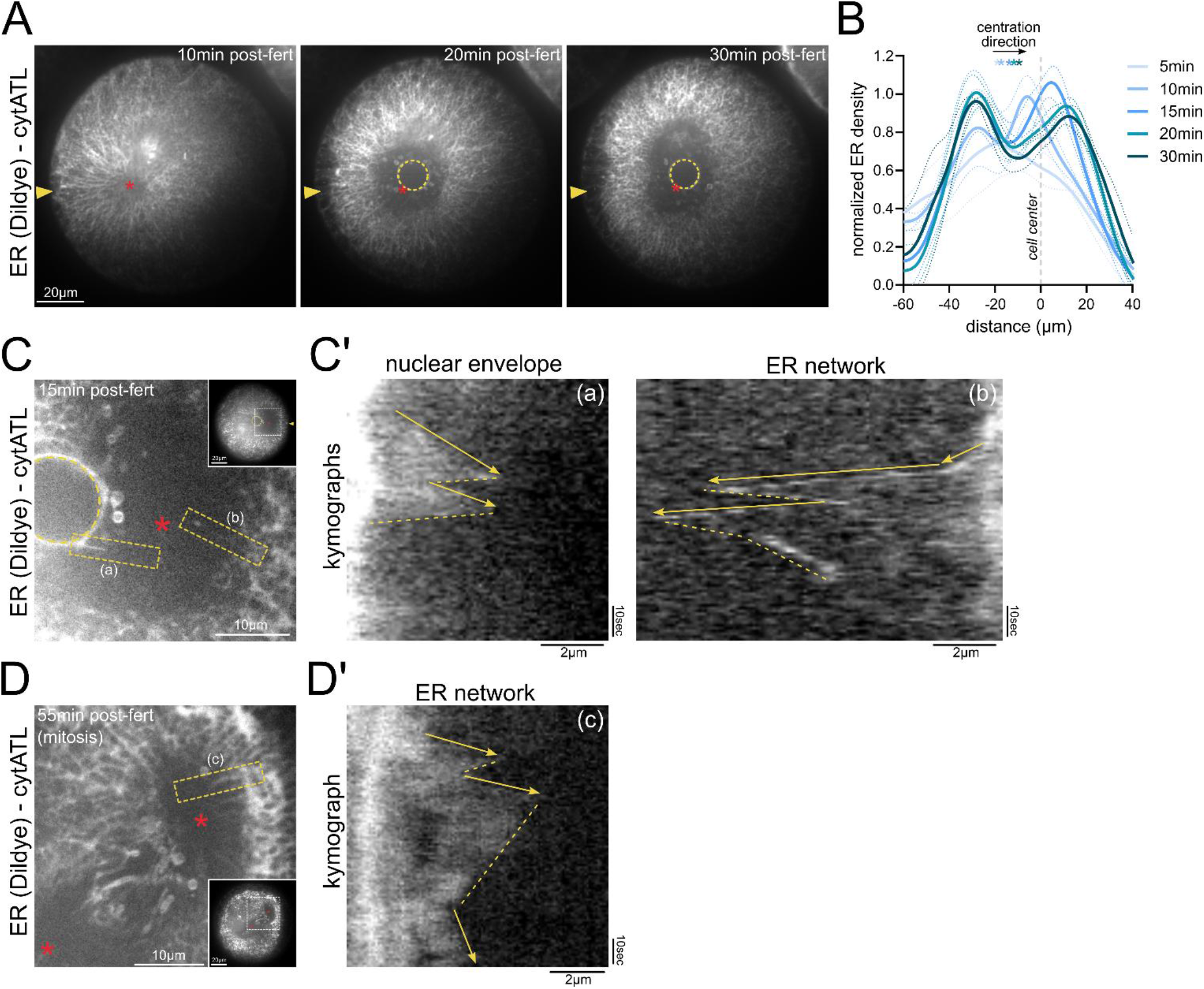
ER fragmentation results from failed ER pulling events. (A) Time-lapse images of aster centration following DiIdye injection (ER labeling) and cytATL protein injection. The yellow arrowhead indicates the fertilization site, the red star marks the aster center, and the yellow dotted circle outlines the female pronucleus. Scale bar, 20 µm. (B) Normalized dynamics of ER density relative to the cell center during aster centration after cytATL protein injection. Each curve corresponds to a specific time point from 5 to 30 min post-fertilization. Dotted lines indicate the standard deviation (SD), and overlaid stars mark the position of the aster center relative to the cell center (n = 4 zygotes). (C-D) Close-up views of single frames from time-lapse images acquired during aster centration (C) and mitosis (D) following DiI dye injection (ER labeling) in zygotes injected with cytATL. Insets show the full zygote view. Red stars mark the position of the aster center or spindle poles. The yellow dotted circle outlines the female pronucleus. Red dotted boxes highlight nuclear envelope and ER membrane pulling events. Scale bar, 10 µm. (C’-D’) Corresponding kymographs of membrane pulling events. Yellow arrows and yellow dotted lines indicate the pulling and retraction phases, respectively.

### A physical model for aster motion based on cytoplasmic ER pulling

In order to test if ER centripetal accumulation may be sufficient to center MT asters, we developed a biophysical model. Recent work suggests that the ER behaves mostly as a fluid, since it can flow and reorganize through membrane fusion and fission events (69). We thus modelled the ER as a fluid of viscosity *η* interwoven with the cytosol (**Fig 7A**). Since we experimentally observed changes in ER density during aster centration, we assumed the ER fluid to be compressible, and implemented repulsions between ER tubules by assuming that ER pressure increases linearly with its density. In contrast, the cytosol was modeled as an incompressible fluid of viscosity *η*_CP_. Moreover, we posited that the motion of the ER with respect to the cytosol results in a frictional drag, with a drag coefficient per unit volume of cytosol *γ* which mechanically couples ER and cytosol. The ER is known to embed dynein motor complexes that can bind to, and walk along MTs, thereby altering ER morphology and density (19,70–72). For simplicity, we considered dynein complexes to be composed of an ER-embedded part and a cytoplasmic part. When the cytoplasmic part binds the embedded part, the complex becomes active, and binds a MT with a given on-rate and move towards its minus end. Here, we assume that the abundance of the cytoplasmic part of the complex is rate-limiting. From this assumption follows that local MT densities, such as that observed in the aster tail, do not impact the model’s output, in agreement with previous works (2,73) (see Methods). Another consequence of this assumption is that we can model only the embedded part of the motor complex, thereafter referred to as the *embedded motor complex* (see Methods). We simplified the geometry of asters by assuming that astral MTs span the whole cell volume. When the aster is off-center, this creates an asymmetry in the aster shape (**Fig 7B**). This asymmetry is more pronounced in the model at early time points but quickly converges to the measured experimental values. In the model, unbound embedded motor complexes are transported by the ER flow, and diffuse in the ER. This diffusion prevents motors from accumulating faster than the ER at the aster center. In contrast, bound embedded dynein complexes move along MTs with a velocity ***v****_m_* towards the aster center. As a result, dyneins exert a force ***f*** = *ζ*(***v****_m_* − ***v***) on the ER, where ***v*** is the local ER velocity, and *ζ* is the friction coefficient of embedded motors in the ER membrane (**Fig 7B**). At the cellular scale, force balance yields two coupled hydrodynamic equations, one for the ER velocity and one for cytoplasm velocity, as well as an equation for aster centration motion (see Methods). Additionally, we write two conservation equations, for the ER and the embedded motor densities, describing their convection and diffusion. Long standing ER density gradients observed experimentally suggest that ER diffusion is small, so that at steady-state the ER flux should vanish. In the limit where ER diffusion is small, dynein forces on the ER are balanced by ER pressure, which is proportional to ER density. Assuming that the diffusion coefficient of embedded dynein complexes in the ER to be large, their density should be proportional to ER density. At steady state, force balance therefore predicted an exponential decrease of ER density away from the aster center (see Methods). Indeed, both EM and live imaging of the aster 30 minutes after fertilization yielded an exponential distribution of ER density. In addition to validating our hypothesis, this yielded a length-scale λ = 30µm, that corresponds to the balance between motor force and pressure in the ER (**Fig 7C**). To study the dynamical behaviour predicted by our model, we numerically solved the model equations, using the FEniCS Finite Element Method (FEM) software (74,75). We chose model parameters (see Methods), to accurately recapitulate experimentally measured aster centration dynamics (8,26) (**Fig 7E**) as well as the ER density length-scale (30 µm, **Fig 7C**) and the timescale for ER packing (10 minutes) (**Fig 3B**, **Fig 7D and 7F**). Lastly, by fitting the ER drag γ, we could reproduce cytoplasmic flow fields with velocities in the range of 0.01 µm/s (**Fig 7G**), similar to that found experimentally (**Fig 2B**). Therefore, this intracellular fluid mechanics model supports the hypothesis that centripetal ER accumulation drives aster motion.

**Figure 7:**
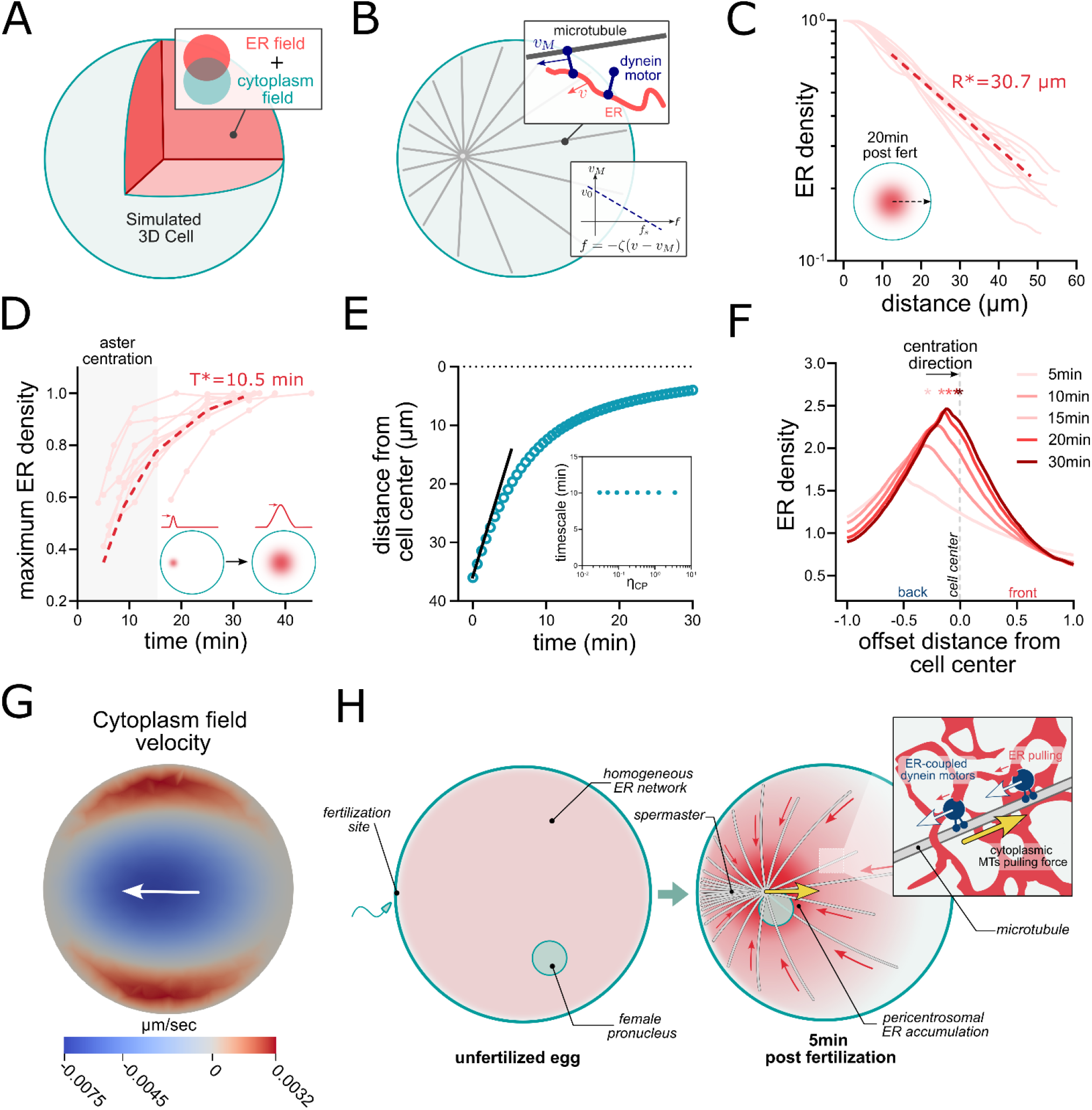
A two-fluid model recapitulates ER accumulation and aster centration. (A) Schematic representing the simulated 3D cell composed of an ER field (compressible fluid) merged with a cytoplasm field (incompressible fluid). (B) Schematic depicting how the simulated aster interacts with ER-bound dynein motors and how the motor velocity-force relationship is modeled with a motor friction term *ζ* (C) ER density profile as a function of distance from the cell center. Light red lines correspond to experimental data. The dashed red line corresponds to the fit used to determine the characteristic ER density length scale R* = 30.7 µm. (D) Evolution of maximum ER intensity. Light red lines correspond to experimental data. The dashed red line corresponds to the fit used to determine the characteristic ER accumulation timescale T* = 10.5 min. (E) Evolution of the distance between the aster center and the cell center in simulated centrations. (F) Simulated evolution of ER density profiles during aster centration. (G) Map of cytoplasm velocity along the aster movement axis *v_x_* in simulation at time t=1min48s. (H) General overview of the working model for aster centration based on ER transport and MT cytoplasm pulling.

## DISCUSSION

In this study, we provide multiple evidences that proper aster centration at fertilization requires an intact ER membrane network. Force exertion is mediated by ER interaction with long MTs projecting at the aster front into bulk cytoplasm, that drag the ER inward to produce an outward centering force (**Fig 7H**). Accordingly, we observed in this region, centripetal cytoplasmic flows and captured stochastic ER centripetal displacement events, close to typical dynein speed and persistence, occurring concomitantly with a global accumulation of the ER at the aster center. Therefore, we propose that the centripetal ER accumulation, and the resulting net drag force applied on the aster, may be sufficient to drive aster centration. An important support of this model is given by experimental manipulations of ER density, which reroutes aster centration (**Fig 4C**) and of ER integrity which halts centration (**Fig 5E-G**). Recent reports have supported a similar role for proper ER/dynein distribution for centrosome positioning in fish and ascidian embryos (17,18). Therefore, our work adds to a growing appreciation for the role of ER transport on MTs as an important force-generating system to position asters in early embryos. In contrast to previous findings in other systems (23,32), we found that the displacement of small compartments like endosomes or mitochondria could only produce forces in the range of tens of pNs, while our previous work allowed to compute a maximal aster centering force on the order of ∼600 pN (25). This is because the typical friction force on a vesicle of radius of ∼200 nm travelling at the dynein maximum speed is on the order of few pNs, implying a constant asymmetry of at least 400 more vesicles on the far side of the aster, to account for values of aster centering force, which we do not observe. Rather, we propose and confirm with our 2-fluid model that the ER acts as a much better cytoplasmic anchor for the aster, because both its internal viscosity and frictional interaction with the cytosol dampen its compaction to optimize force exertion. With our model parameters, both effects have a similar order of magnitude since we find the normalized ER drag (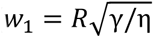) to be 2. This ER drag is in the range expected from simple scaling arguments (see Methods), and corresponds to a total ER drag *γ V* ∼ 0.05 s N/m, with *V* the total cell volume, around five times larger than the previously measured for aster drag (25). A consequence of this large frictional interaction and internal ER viscosity, is that in our model, motors move at speeds close to that of the aster, but much slower than the typical unloaded speed of dynein motors, *v_MOT_∼v_aster_∼v_0_*/10 (see Methods), so they exert a force close to their maximal stall force (Toba et al., 2006). In contrast, when motors share the load of moving a single vesicle, they may move faster and exert much lower forces. As such, the few discernable events of fast ER movements observed in our experiments (**Fig 6C-D**) may correspond to local weakening of the ER structure, that allow motors and ER tubes to move fast but exert non-productive small forces to the aster. Accordingly, the high occurrence of these unproductive pulling events observed when cytATL is injected reflects the more brittle state of the ER, and leads to a general abolishment of MT/motor force exertion and consequent aster stalling. In conclusion, pulling on a strong anchor that accumulates slowly allow motors to exert larger forces, implicating a smaller number of motors in the large ∼600 pN force required to center the aster.

The status of other force-generation mechanisms and especially whether they compete and/or cooperate with ER-based cytoplasm pulling, remain to be fully elucidated. Their relative contributions, influenced by factors such as cytoplasm organization, cortical polarity and cell geometry, may account for the diversity of asters and spindles positioning phenotypes and consequent species-specific architecture of cleavages of early embryos of most metazoans (1,76–78). In sea urchins, a brief burst of F-actin polymerization at the fertilization site appears to provide an initial push to the aster. However, when this process is disrupted, aster displacement is largely compensated by MT-dependent forces. This two-step mechanism closely resembles what has been described in the mouse zygote, although in sea urchins it occurs within a much shorter time frame (47). MT-based cortical pushing has also been proposed as an important alternative mechanism and was suggested to represent the dominant mechanism for aster centration (8). A key experiment supporting this hypothesis relied on the use of the p150^CC1^ fragment to inhibit dynein activity. However, our complementary experiments indicate that dynein inhibition by p150^CC1^ is only partial, during the short time-scale of aster centration, and produces a broad spectrum of altered aster trajectories. Among these, interrupted motions are consistent with progressive inhibition of dynein motor activity and loss of cytoplasmic pulling forces. Yet, it remains unclear why in some cases the aster moves toward the opposite side of the zygote or initiates spiral-like trajectories. Interestingly, similar large-scale movements were also reported in encapsulated Xenopus egg extracts following dynein inhibition (79), and were proposed to depend on the formation of MT bundles and kinesin motor activity, mediated by MT–MT sliding, rather than MT polymerization. By analogy, these late phenotypes observed after p150^CC1^ treatment in sea urchin zygotes could reflect a global reorganization of the MT network, favoring kinesin-based forces in the context of weakened dynein activity. Developing fast and highly specific methods for dynein inhibition will likely be essential to fully disentangle the relative contributions of cytoplasmic pulling versus cortical pushing forces for aster positioning.

The role of the endoplasmic reticulum (ER) as a cytoplasmic anchor for aster motion implies the existence of a molecular linkage between MTs, dynein complexes, and ER membrane proteins that enables the transmission of ER drag forces to MTs. Although minus end–directed ER pulling events similar to those described in this study have been reported in multiple systems, including Xenopus eggs, Drosophila spermatocytes, and cultured vertebrate cells (80–82), the molecular nature of this coupling still remain largely elusive. In the specific context of ER exit sites, however, dynactin was shown to interact with the ER coat protein Sec23p to promote directional transport toward the Golgi apparatus (83). More recently, CDR2 has been identified as a novel ER adaptor for cytoplasmic dynein. This interaction, mediated by the ER membrane protein kinectin (KTN1), appears to regulate the subcellular distribution of ER sheets (70). In addition to ER–dynein coupling, MTs have also been shown to interact directly with ER proteins such as Climp63, REEP1, p180, or KTN1 (84). These interactions are consistent with the fact that ER and MTs are found interlaced in many cellular contexts which may here contribute to stabilize the growing aster structure and also prevent fast ER re-homogenization between pulling events. In a similar fashion, EB1-mediated interaction with the ER protein STIM1 described in cultured cells may facilitate microtubule growth in the crowded egg cytoplasm (85). Thus, ER-aster interactions are likely more complex than what depicted here and identifying specific adaptor proteins responsible for aster centration may represent an important step toward elucidating the molecular mechanisms underlying MT force generation in the cytoplasm.

## MATERIALS AND METHODS

### Sea Urchins

Adult purple sea urchins (*P. lividus*) were shipped from the Roscoff Marine station (France) and kept at 16°C in an aquarium for several weeks in ASW (Artificial Sea Water - Reef Crystals, Instant Ocean). Gametes were collected by intracoelomic injection of 0.5 M KCl. Sperm was collected dry and kept at 4°C for up to 1 week. Eggs were rinsed twice, kept at 16°C, and used on the day of collection.

### Microscopy

High-resolution microtubule imaging of fixed and live samples was performed using an LSM980 AiryScan2 confocal microscope (Zeiss) equipped with a 60× water-immersion objective (NA 1.2) and operated in SR-4Y mode. The microscope was controlled using ZEN software (Zeiss). Live imaging of the endoplasmic reticulum, spherical cytoplasmic vesicles, and microtubule plus ends was performed on a spinning-disk confocal microscope (TI-Eclipse, Nikon) equipped with a CSU-X1FW confocal scanner unit (Yokogawa), an ImageEM-1K EM-CCD camera (Hamamatsu) or a Prime BSI sCMOS camera (Teledyne Photometrics), and either a 40× (NA 1.25) or 60× (NA 1.42) water-immersion objective. This system was operated using MetaMorph® software (Molecular Devices). Aster trajectories were imaged using a widefield fluorescence microscope (TI-Eclipse, Nikon) equipped with a 20× dry objective (NA 0.75) and an ORCA-Flash4.0 LT CMOS camera (Hamamatsu), operated with Micro-Manager software (Open Imaging).

### Immunostaining

Before fertilization, eggs were treated with 0.01 mg/mL trypsin in ASW for 10 min, followed by two rapid washes in fresh ASW to remove the vitelline envelope. Eggs were then fertilized, and zygotes were fixed 5 min after fertilization in fixation buffer (100 mM HEPES, pH 6.9; 50 mM EGTA; 10 mM MgSO₄; 400 mM dextrose; 0.2% Triton X-100; 0.2% glutaraldehyde; 2% formaldehyde) for 1 h. Samples were subsequently washed in 0.1% Triton X-100 in PBS, transferred to fresh PBS, and incubated in 0.1% NaBH₄ in PBS for 30 min at room temperature. After a brief PBS wash, samples were returned to 0.1% Triton X-100 in PBS and incubated overnight at 4°C with a mouse anti-tubulin antibody (DM1A, ThermoFisher) diluted 1:5000 in 0.1% Triton X-100 in PBS. The primary antibody was washed extensively (>1 h) with three washes in 0.1% Triton X-100 in PBS at room temperature. Samples were then incubated overnight at 4°C with an anti-mouse IgG1 secondary nanobody (Nano-Secondaries™) conjugated to Alexa Fluor™ 488, together with phalloidin CF®568 (Biotium), diluted to 1:1000 and 0.2 U/mL, respectively, in 0.1% Triton X-100 in PBS. Finally, samples were washed three times in 0.1% Triton X-100 in PBS, incubated with 10 µg/mL Hoechst 33342 (Thermo Fisher Scientific) in PBS for 15 min, and mounted in mounting medium (90% glycerol, 0.5% n-propyl gallate in PBS).

### Microinjection

The jelly coat surrounding unfertilized eggs was mechanically removed by passing them three times through an 80µm Nitex mesh to allow them to adhere on protamine-coated glass bottom dishes (MatTek Corporation). Once transferred on the protamine-coated dish, eggs remained unfertilized for maximum of 15 min before microinjection and fertilization. Injection needles were prepared from borosilicate glass capillaries (1 mm diameter) using a needle puller (P-1000; Sutter Instruments) and a micro-grinder (EG-400; Narishige). Microinjections were performed using a motorized micromanipulation setup (FemtoJet 4, Injectman 4; Eppendorf) mounted on a widefield fluorescence microscope (see above). Protein injections were performed such that the injected volume corresponded to 1–5% of the total egg volume. The estimated final intracellular protein concentrations were 20–100 µg/ml for p150^CC1^ and *pliv*p150^CC1^, 4-20 µg/ml for TMBD-mStayGold, 10-50 µg/ml for *pliv*EB1-GFP, cytATL, and cytATL^R232Q^, 0.5-2.5 µg/ml for H2A-H2B-Chromobody-mScarlett3 and 2-10 µg/ml for UtrCH. Live endoplasmic reticulum (ER) labeling was achieved by injection of a soybean oil droplet saturated with a lipophilic fluorescent dye (DiIC18(3); Thermo Fisher) (57). When combined with protein injection, oil injection was performed first, followed by protein injection after a 5–10 min recovery period.

### Egg centrifugation

Cytoplasm stratification approach was based on centrifugation protocols from previous studies using various echinoderm species (86–88). In brief, non-fertilized eggs in ASW were carefully deposited on the top of an equal volume of 0.85M sucrose solution in a microtube. The tube was then gently tilted to establish a density gradient between the two solutions and centrifuged for 7 to 10min at 10000g. Stratified eggs were quickly recovered by pipetting few microliters where they equilibrated in the density gradient, briefly washed in fresh ASW and used immediately for experiments.

### “Stalled aster” assay and cytoplasmic flow analysis

Previously described magnetic tweezers were used to apply a force on the centered aster and displace its center toward cell cortex (89). In brief, right after centration completion, zygotes containing injected magnetic particles were approached from a magnetic tip until magnetic particles and aster center were pulled back toward the cell cortex. The aster was held in position for few minutes to ensure residual flows due to aster displacement were dissipated before recording DIC time-lapses at a rate of 1, 2 or 3 images per seconds. Flow analysis has been performed using the particle image velocimetry PIVlab tool in MATLAB. Contrast-limited adaptive histogram equalization and two-dimensional Wiener filter with accordingly windows of 20- and 3-pixel widths were applied to the images in the pre-processing steps for noise reduction. Image sequences were investigated in the Fourier space by three interrogation windows with 64-, 32-, and 16-pixels widths and 50% overlapped area. The spline method was used for the window deformation and subpixel resolution obtained by two-dimensional Gaussian fits. The distribution of velocity components of vectors for each set was visually inspected and restricted to remove outliers in the post-processing stage. Moreover, two other filters based on the SD and local median of velocity vectors were applied to validate the vector fields. The output vector fields after smoothing were used for plotting flow maps and vector fields at different time intervals.

### Aster center and microtubule plus ends tracking

Growing microtubule plus ends were labeled *via* the injection of EB1–GFP protein. The male pronucleus was labeled with 5 µg/mL Hoechst 33342 in ASW and used as a proxy for centrosome position; following pronuclear fusion, the zygote nuclear center was used as a proxy. EB1 comets and male pronuclei were tracked manually using the TrackMate plugin in Fiji. Aster motion was subtracted from EB1 comet trajectories using a custom-written Python script to compute polymerization velocities relative to the aster center.

### Cytoplasmic vesicles tracking and cytoplasmic pulling force estimation

Vesicular compartments were labelled using commercially available specific fluorescent probes. Unfertilized eggs were incubated for 30-45 min with Nile Blue (1:2000, Sigma-Aldrich®), FM1-43 (1:1000, ThermoFisher®), LysoTracker (1:1000, ThermoFisher®), or MitoTracker (1:500, ThermoFisher®) to respectively label yolk granules, endosomes, lysosomes, or mitochondria. Individual vesicles were automatically tracked using the TrackMate Fiji plugin (90). In parallel, the aster center was manually tracked using Hoechst signal (male pronucleus). The analysis of the trajectories was performed using a custom-written Matlab script. Trajectories were initially filtered to be longer than three seconds therefore consisting of at least four time points. To approximate the total force exerted on a microtubule aster, the exerted drag forces of all vesicles along the aster displacement axis in a rotated coordinate system were calculated. The weighted average force was obtained by the weights that were the net portions of the trajectories that the vesicles were moving inward/outward of the aster considering the moving directions. The Stokes’ drag was calculated as f=6πηr∑vx where η was the viscosity of cytoplasm and equal to 1000X water viscosity and r was the size of the vesicles. The vesicle sizes were based on SBF-SEM data. The summation is over the x-component of instantaneous velocity.

### Kymographs

Rapid local movements of the ER membrane network were analyzed using the Dynamic Kymograph plugin in Fiji. Kymographs were generated along deformable lines oriented radially with respect to the aster center, allowing for compensation of aster displacement and cytoplasmic background noise. Duration, amplitude and velocity of ER pulling events were computed from kymographs

### Drug treatments

Nocodazole treatment was applied 1 to 2 minutes prior fertilization at a final concentration of 20µM. Latrunculin B treatment was applied immediately after the visual confirmation of fertilization at a final concentration of 20µM. Both chemicals were prepared from a 2mM stock solution (100X) in DMSO and pre-mixed with 1ml of ASW before being added to the culture dish.

### Electron Microscopy & Cytoplasm Segmentation

Details of endomembranes organization have been captured using serial block face scanning electron microscopy (SBF/SEM). Unfertilized eggs and Zygotes were fixed in 0.2M sodium cacodylate, 0.25M sucrose, 2% paraformaldehyde, 2% glutaraldehyde for 1h at room temperature. Fixed sample were then progressively transferred in 0.2M sodium cacodylate, 0.35M NaCl. Samples were further processed for SBF/SEM following the NCMIR protocol (https://ncmir.ucsd.edu/sbem-protocol) as previously described (51). Processed samples were mounted on aluminum pins, trimmed and inserted into a TeneoVS SEM (Thermo Fisher Scientific). Acquisitions were performed with a beam energy of 2.7kV, a current of 400pA, in LowVac mode at 40Pa, a dwell time of 1μs per pixel and sections of 100nm (voxel size 20x20x100nm). Yolk granules, mitochondria and endoplasmic reticulum were segmented from the resulting image stacks with a supervised learning approach using the Ilastik software (1.4.0-gpu) (91). Pixel classifications were obtained through an independent autocontext workflow for each compartment. 3D renders were generated from the pixel probability maps in Blender using the *microscopynodes* add-on (92).

### Plasmids and Protein purification

TMBD-mStayGold (Tau Microtubule Binding Domain) construct has been derived from TMBD-GFP (41). Plasmids used to express cytATL and cytATL^R232Q^ proteins were obtained from the Tom Rapoport laboratory (63). Plasmids used to express the Utrophin CH domain (UtrCH) and H2A-H2B-Chromobody-mScarlett3 were obtained from the Peter Lenart laboratory (93). Purified UtrCH protein was subsequently labelled using CF633™ maleimide following manufacturer’s instructions (Sigma-Aldrich®). Plasmid used to express p150^CC1^ protein (44) was obtained from the Jesse Gatlin laboratory. An alternative version of this construct has been generated using the p150^CC1^ sequence form *P. lividus* (Eurofins Gene Synthesis) which was subcloned in a pRSET-A vector. The EB1 sequence from *P. lividus* has been obtained through gene synthesis (Proteogenics) and subcloned in a pET22b vector for EB1-GFP protein expression. All recombinant proteins were expressed in BL21 (DE3) *E. coli* strain (ThermoFisher®) and purified using either EZview™ Red HIS-Select® HC Nickel Affinity Gel (Millipore®) for TMBD-mStayGold, UtrCH, *pliv*EB1-GFP and *pliv*p150^CC1^ proteins; or EZview™ Red Glutathione Affinity Gel (Millipore®) for p150^CC1^, cytATL and cytATL^R232Q^ proteins. When required, proteins were concentrated using 10-30KDa MWCO Amicon® centrifugal units (Millipore®).

### Physical model of the ER

#### Geometry and physical assumptions

For the sake of simplicity, we consider that the cell is spherical and that MTs form an aster with a center initially located away from the cell center. MTs are uniformly distributed radially around the aster center and extend until their + ends touch the cell cortex. During centration, the aster center moves toward the cell center, and the aster keeps filling the cell volume The aster is thus highly asymmetric at the start, and becomes more and more symmetric as it centers. Aster motion is due to motors anchored to the ER that are bound to MTs. In the case of dyneins, it is known that active motors are complexes that include the dyneins themselves, dynactin and activating cargo adaptors (94–96). Here, we assume that active motors can bind and unbind MTs; when bound to MTs, they walk toward the aster center at a velocity **v***_m_* and create forces that pull on the aster. Because an off-centered aster breaks front-rear symmetry, the spatial distribution of force-generating motors is asymmetric, and the resulting net force pulls the aster center toward the cell center, driving the centering motion.

#### Motor complex transport and binding kinetics

We consider the transport of the motor complex, of which part is embedded in the ER, and part is diffusing or actively transported in the cytosol. Because of the intricacies of motor activation, we chose to use a simplified model for motor complex dynamics. Here, we assume that some of the cytoplasmic part of the complex (for example, an adaptor) is the rate-limiting element. In this case, we may consider explicitly only the fraction of the complex embedded in the ER, while the cytoplasmic part will implicitly control the magnitude of the binding rate of unbound motors to MTs. In the following the term “embedded motor complex” will be used as a shorthand for the membrane-bound part of the complex. Unbound embedded motor complexes at concentration *c_m_^u^* are carried by the ER and can diffuse relative to the ER in which they are embedded. The ER structure is very heterogeneous and difficult to characterize precisely. We therefore describe it as an isotropic compressible fluid. In this description, motor diffusion is measured relative to the ER material itself rather than to the reference frame of the cytosol. Thus, zones of higher ER density will tend to embed more motor complexes, which gives rise to an ER concentration-dependent effective embedded motor diffusion. Taking *ρ*_0_ as a reference ER concentration, we assume that local ER compression and dilution are isotropic. Carefully considering diffusion in the Lagrangian frame shows that the concentration of unbound embedded motor complexes is driven by the concentration of the ER through the conservation equation:

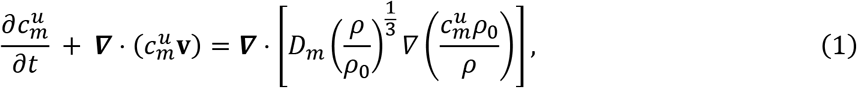

where **v** denotes the local velocity of the ER.

The corresponding current of unbound embedded motor complexes is:

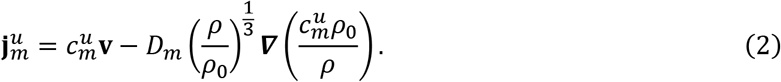

This form shows that diffusion relaxes the gradients of motor concentration measured per unit ER material, 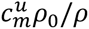.

Bound embedded motor complexes walk on MTs, but their velocity is not the free motor velocity because they are slowed down by the friction force exerted by the ER on the motor:

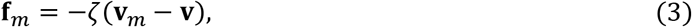

where *ζ* is the friction coefficient between ER and motors, that may depend in a complex way on the molecular architecture of the ER. The local orientation of MTs defines a unit vector **u** toward the aster center, and the velocity of the motors relative to the MTs is along **u**. This velocity is given by the force-velocity relation, which we approximate by the linear relation (97):

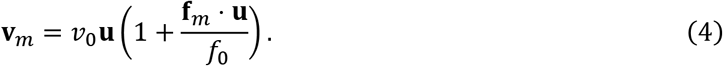

Here, *f*_0_ is the stall force of the motor and *v*_0_ is its velocity at vanishing load. This results in the motors slowing down when a force opposes their movement, a consequence of thermodynamics limitations. When numerically solving our sets of equations, we find that motors move at a velocity close to the aster velocity, much smaller than *v*_0_. This means that they exert a force close to their stall force *f*_0_.

In the following we introduce the dimensionless quantity 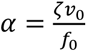. The current of bound motors is

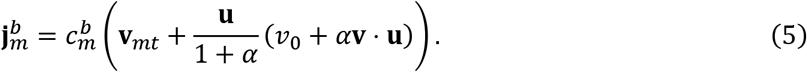

We also need to describe the binding and unbinding of embedded motor complexes to MTs. We assume unbound embedded motor complexes to bind reversibly to the (rate-limiting) cytoplasmic factor c_cyto_, and the resulting complex can then bind MTs:

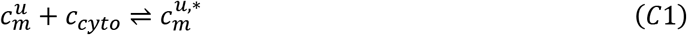

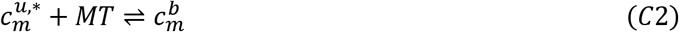

We also assume that the binding and unbinding kinetics are fast compared to the flow, such that the binding reaction is at local equilibrium, i.e. 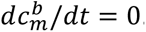. This yields 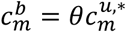 and following Michaelis-Menten kinetics, we find the generic formula:

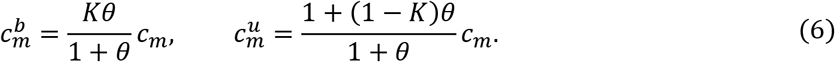

We defined *θ*(**r**) = *ρ_mt_*(**r**)/*ρ_sat_* the dimensionless MT density, with *ρ_mt_*(**r**) the MT density and *ρ_sat_* a characteristic density (defined as the ratio of the off-rate to the on-rate in reaction C2). K is a dimensionless control parameter, and both K and *ρ_sat_* depend on the reaction rates and availability of the cytoplasmic part of the dynein complex *c_cyto_*. Experimentally, we find that the aster movement seems mostly independent of MT density, since the aster does not move backwards in spite of its the dense MT tail. This hints that MTs are not the limiting factor to motor activity, in agreement with our assumption that motor activity is controlled by the cytoplasmic part. We can therefore take the limit *θ*(**r**) → +∞, and we find 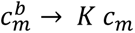 and 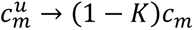, i.e. the density of bound and unbound embedded complexes is independent of MT density. In that limit, K becomes the fraction of embedded motor complexes bound to MTs. If the diffusion constant of the unbound embedded motor complexes is very large, we have 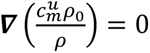. In this limit, the concentration of unbound embedded motor complexes is proportional to the ER concentration 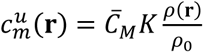. With 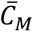 the mean concentration of embedded motor complexes. Eventually, the total concentration of embedded motor complexes *c_m_* is a conserved variable and 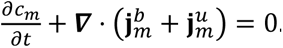.

#### ER and cytoplasm hydrodynamics

We consider the ER as a compressible viscous fluid with a shear viscosity *η* and a bulk viscosity *ξ*. The mass conservation equation of the ER is

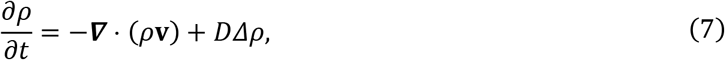

where *D* is the ER diffusion constant.

The ER interacts both with the molecular motors and with the cytoplasm. The force exerted by one motor on the ER is

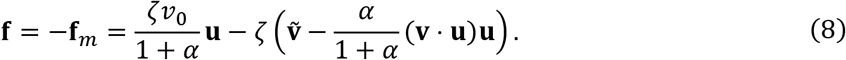

The cytoplasm is an incompressible viscous fluid with viscosity *η*_cp_. In order to take the interactions between the ER and the cytoplasm into account, we use a two-fluid model with a friction coefficient *γ*. The equation for the ER velocity is

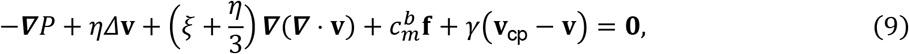

where *P* is the pressure of the ER fluid, which depends on its concentration *ρ*, and **v**_cp_ is the local velocity of the cytoplasm. The equation for the cytoplasmic velocity is

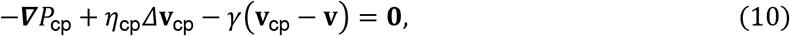

where *P*_cp_ is the pressure in the cytoplasmic fluid, which can be considered as a Lagrange multiplier ensuring the incompressibility condition ***∇*** ⋅ **v**_cp_ = 0.

#### Aster force balance and motor-limited closure

The force exerted by one motor on the aster is equal to the friction force on the motor due to the ER, **f***_m_*, because the total external force on each motor must vanish. The total force exerted by the motors on the aster is balanced by the friction force of the aster on the cytosol. If the position of the aster is **r**_0_, the friction force is (1/*μ*)*∂_t_***r**_0_, where *μ* is the aster mobility. The velocity of the aster is therefore

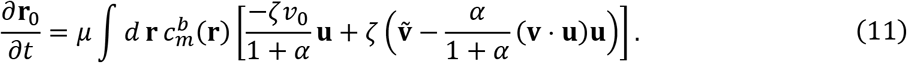

where **u** = (**r**_0_ − **r**)/|**r**_0_ − **r**|

Because the Stokes equations (9) and (10) are linear, the velocity field **ṽ** can be decomposed into a part generated by the motor forces for a fixed aster and a part proportional to the translational velocity of the aster, **v***_mt_* = *∂_t_***r**_0_. When inserted into Eq. (11) the contribution proportional to **v***_mt_* gives a force proportional to *∂_t_***r**_0_. It therefore acts as an additional friction on the moving aster, rather than as an independent active centering force. Since the bare aster mobility *μ* is not known independently, this motor-induced friction can be absorbed into an effective phenomenological mobility. In the following we therefore compute the active driving force in the frame of the aster, setting **v***_mt_* = 0 in the force-generating terms.

#### Steady-state ER density

Assuming the diffusion to be near-zero, steady-state implies **v** = **v**_cp_ = 0. Eqn. (9) then leads to 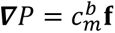. We assume unbound embedded motor complex diffusion to be fast in the ER, such that:

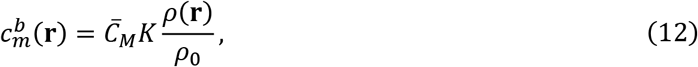

With 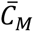 the mean concentration of embedded motor complexes. Taking Eq. (12) and a pressure density relation *P* = *Aρ* we obtain a steady-state density profile

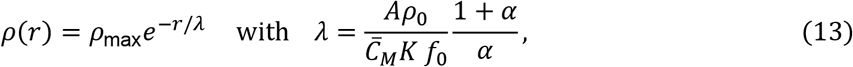

with *r* the distance from the cell center and *ρ*_max_ the ER density at the aster center.

#### Numerics

We solve the ER and motor conservation equations together with the hydrodynamic equations for the ER and cytoplasm using a continuous Galerkin finite-element discretization implemented with the python interface for FEniCS (74,75), on three-dimensional spherical meshes generated with Gmsh (98). The scheme is semi-implicit in time: at each time step, all field equations are solved implicitly, whereas the aster position is updated explicitly from the motor forces computed at that step. Parameter estimation showed that the dynamics are mainly controlled by four sensitive parameters, which were fitted to the centration timescale, the ER packing length scale and timescale, and the magnitude of cytoplasmic flows. Other parameters, such as *α*, *D*, and *D_m_*, lie in limiting dimensionless regimes and only weakly affect the numerical results.

**Table 1:** Values of known parameters used in the model.

| Parameter | definition | value | source |
| --- | --- | --- | --- |
| $\mu$ | Aster mobility | 100 m s<sup>-1</sup> N<sup>-1</sup> | (25) |
| $\eta_{cp}$ | Cytoplasm viscosity | 500 mPa.s | (52) |
| $v_0$ | Dynein unloaded speed | 0.75 $\mu\text{m}$ / s | (99) |
| $R$ | Cell radius | 45 $\mu\text{m}$ | (25) |

**Table 2:** Values of sensitive unknown parameters and combinations used in the model and estimated through fitting.

| Parameter | definition | Fitted value | Strongly impacts |
| --- | --- | --- | --- |
| $w_2 = R\sqrt{\bar{C}_M K / \eta(1 + \alpha)}$ | Relative pull | 0.93 | All observables |
| $w_1 = R\sqrt{\gamma / \eta}$ | Relative drag | 2.0 | Cytoplasm velocity |
| $A/\eta R^2 v_0$ | ER packing pressure | 0.44 | ER packing length scale |
| $\eta/\eta_{cp}$ | Relative ER viscosity | 13.33 | All observables |

**Table 3:** Values of other parameters, to which model results are not very sensitive.

|  |  |  |
| --- | --- | --- |
| $D/Rv_0$ | Effective ER diffusion | 0.001 (small) |
| $D_M/Rv_0$ | Effective motor diffusion | 1 (large) |
| $\alpha = \zeta v_0 / f_s$ | Motor efficiency | 1000 (large) |
| $K = c_m^b / c_m$ | Fraction of bound motors | 0.5 (large) |

### Estimation of ER volumetric drag **γ**

There is no simple estimation of the ER volumetric drag because of its complex morphology. However, we can estimate some bounds. We consider the ER as a network of membrane tubes of 50nm radius, of density 7.5%. From this we can estimate a pore size of approximately 500nm, and a total length of membrane tubes of approximately 6m. Assuming permeability to scale like the pore size square, Darcy’s law on porous media predicts a relative drag 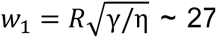. However, this is a gross overestimation, since there are regions where the ER is much less dense, allowing cytosol to flow around, rather than through ER pores. Another, slightly more realistic higher bound for the drag can be computed by estimating the friction of ER tubules. Using our estimation of the total length of ER tubes and the friction of slender bodies, we can estimate *w*_1_∼10. However, this is most likely an overestimation. Even without considering the actual ER structure, this neglects hydrodynamic interactions between nearby tubes, that would lower the effective drag. On the other hand, we can estimate a lower bound for the drag of the ER would have if these hydrodynamic interactions were infinite. In such case, the ER would have the drag of a solid ball of radius R in the cytosol. Neglecting boundary conditions, we can write γ∼6πη_CP_R, yielding *w*_1_∼0.6. In our model, we used *w*_1_∼2, the geometric mean of the lower and upper bounds, to be a good fitting parameter.

## ACKNOWLEDGMENTS

We thank all members of the Minc team for discussion and technical help. We acknowledge the ImagoSeine core facility of the Institut Jacques Monod, member of the France BioImaging infrastructure (ANR-24-INBS-0005 FBI BIOGEN) and GIS-IBiSA and the support of the Région Île-de-France (Sesame). DW received funding from the European Research Council (ERC) ERC-SyG (Grant agreement ID: 101071793) (awarded to Pierre Sens, Institut Curie). H.P. was supported by a fellowship from the ‘‘Fondation pour la Recherche Médicale’’ (FRM) (FDT201904008226). J.N. was supported by a fellowship from the ARC foundation (PDF20191209818) and was an EMBO Non-Stipendiary Fellow (ALTF 881-2019). This work was supported by the Centre National de la Recherche Scientifique, the Université Paris Cité, and grants from La Ligue Contre le Cancer (EL2021.LNCC/NiM) and the Fondation Bettencourt Schueller (“Impulscience”) to N.M.

## Author contributions

Project design, J.S., S.D. and N.M.; Experiments, J.S, J.N. C.M.D., H.P., I.M., A.G., and N.J.; Electron microscopy, R.L.B., C.D. and J-M. V.; Finite element simulation, S.D., D.W., and J-F. J.; Original Draft, J.S., S.D. and N.M.; Draft editing, J.S., S.D., D.W. J-F.J., and N.M.

## SUPPLEMENTARY FIGURES

**Figure S1:**
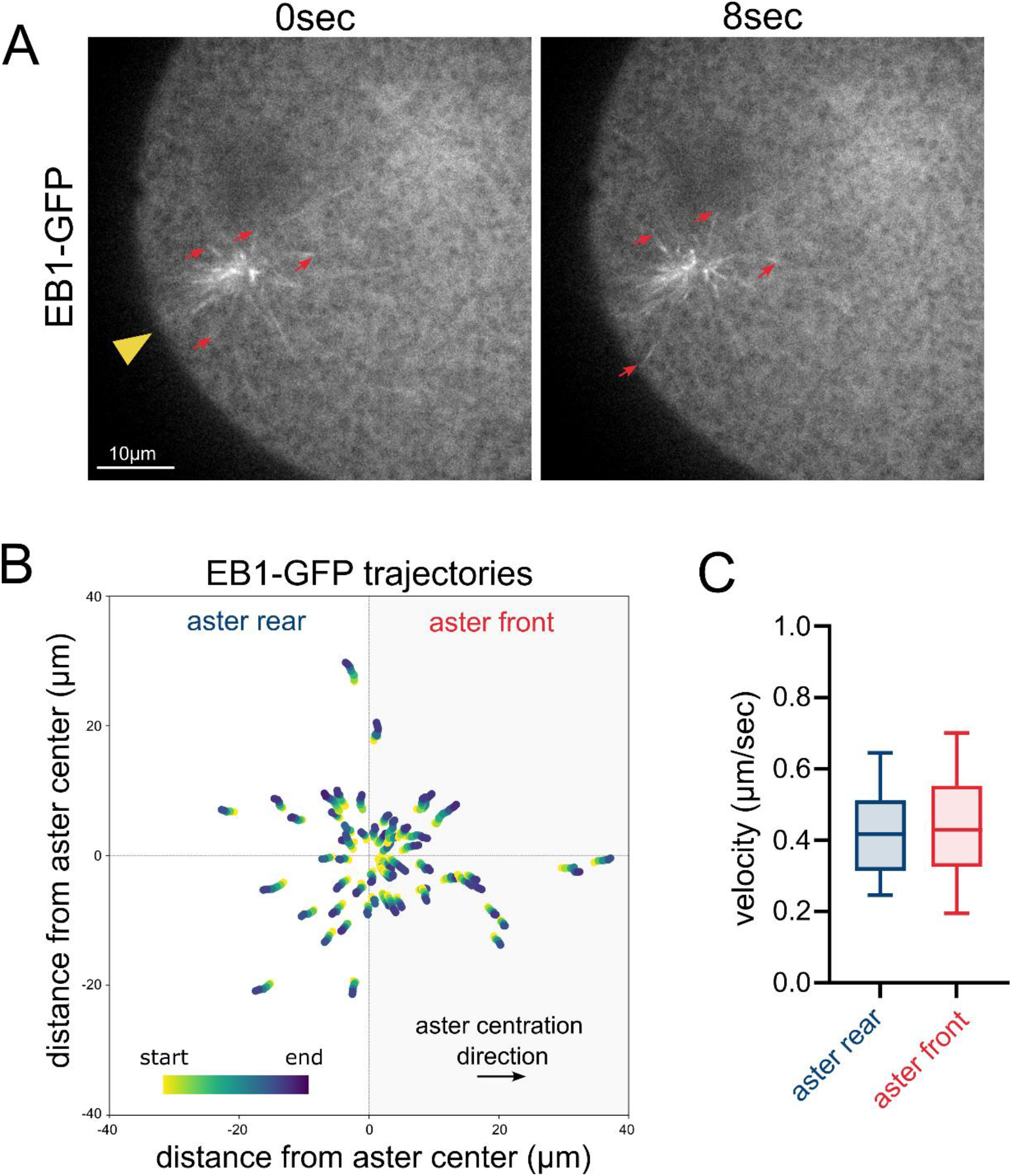
Microtubule plus-end dynamics during aster centration. (A) Time-lapse images of aster centration in a zygote injected with EB1–GFP. The yellow arrowhead indicates the fertilization site, and yellow arrows point at growing microtubule plus ends. Scale bar, 10 µm. (B) Trajectories of EB1 comets plotted relative to the aster center (n = 60 tracks from 3 zygotes). (C) Velocity of EB1 comets in the front and rear of the aster. Error bars represent SDs.

**Figure S2:**
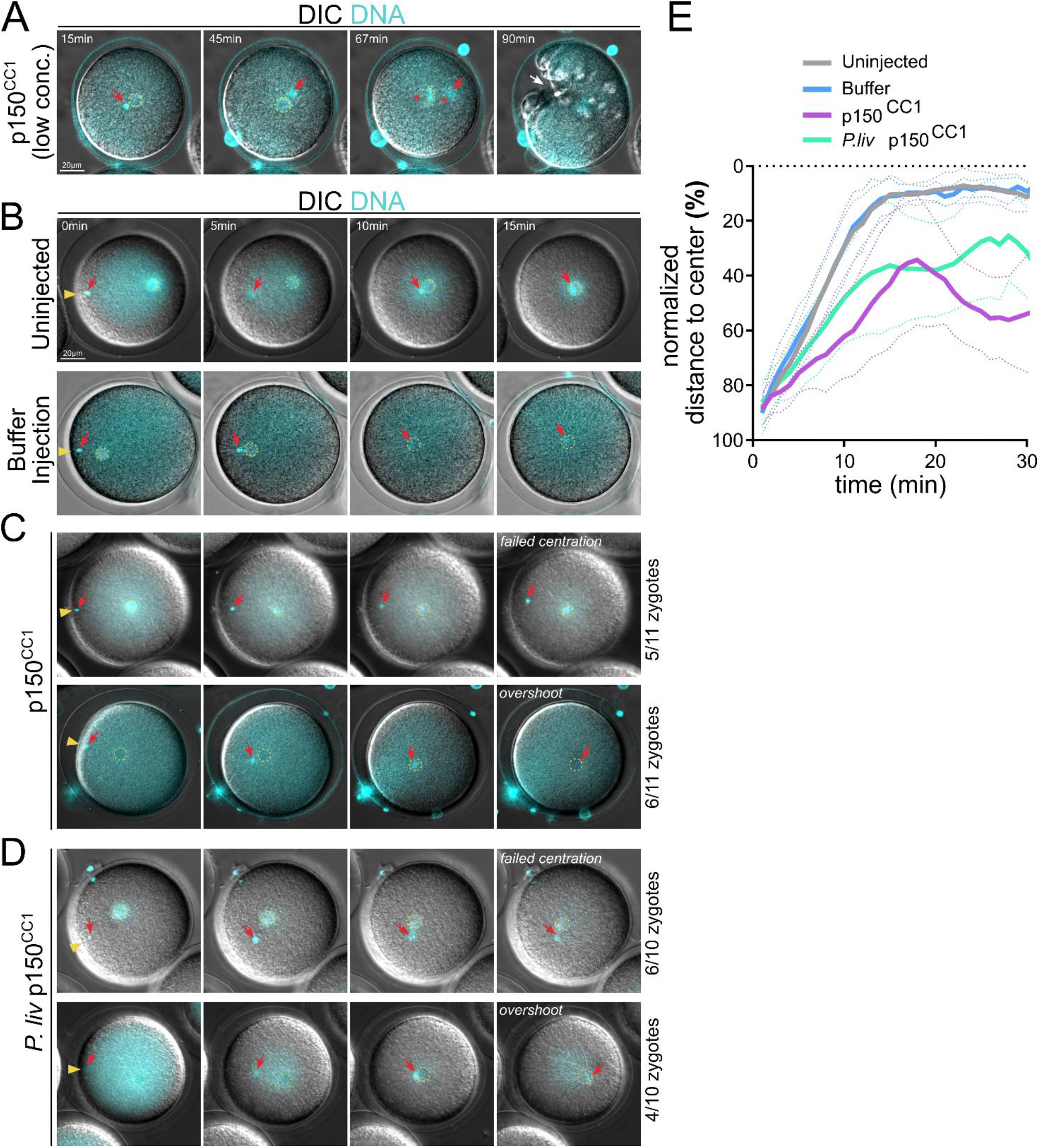
Aster centration defects after p150^CC1^ mediated dynein inhibition. (A-D) Time-lapse images acquired during aster centration. The DIC channel is shown in white and DNA (Hoechst 33342) in cyan. The yellow arrowhead indicates the fertilization site, the red arrow marks the aster center, and the yellow dotted circle outlines the female pronucleus. Scale bar, 20 µm. (A) Post-centration phenotype of the injection of p150^CC1^ at low concentration (≈0.25 µg/ml). The white arrow shows the partial furrow ingression. (B) Non-injected and buffer-injected control zygotes. Representative examples of “failed centration” and “overshoot” aster trajectories in zygotes injected with p150^CC1^ and *P.liv*-p150^CC1^ (≈50-100 µg/ml) are shown in (C) and (D), respectively. (E) Dynamics of the normalized average distance between the aster center and the cell center in non-injected zygotes (gray, n = 8), buffer-injected zygotes (blue, n = 6), and zygotes injected with p150^CC1^ (purple, n = 10) or *P.liv*-p150^CC1^ (green, n = 9). Dotted lines indicate the standard error.

**Figure S3:**
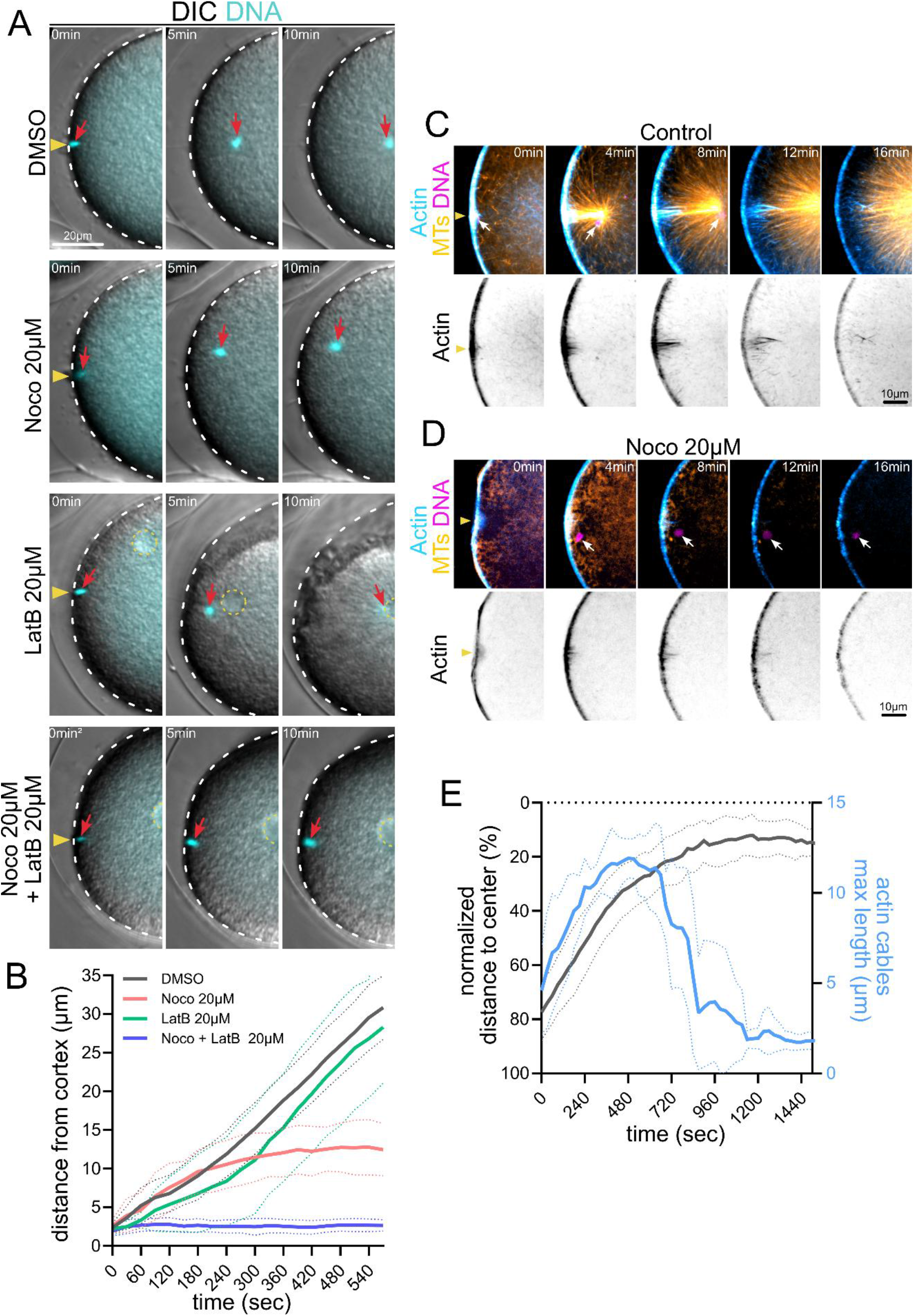
Actin contribution to aster centration is limited. Time-lapse images acquired during early aster centration in zygotes treated, from top to bottom, with DMSO, 20 µM nocodazole, 20 µM latrunculin B, or a combination of 20 µM nocodazole and 20 µM latrunculin B. The DIC channel is shown in white and DNA (Hoechst 33342) in cyan. The yellow arrowhead indicates the fertilization site, the red arrow marks the aster center, the white dotted line outlines the zygote boundary, and the yellow dotted circle outlines the female pronucleus. Scale bar, 20 µm. (B) Dynamics of the average distance between the aster center and the cell cortex in zygotes treated with DMSO (black, n = 14), 20 µM nocodazole (red, n = 12), 20 µM latrunculin B (green, n = 10), or 20 µM nocodazole plus 20 µM latrunculin B (blue, n = 13). Dotted lines indicate the standard error. (C-D) Time-lapse images of early aster centration in zygotes injected with TMBD–mStayGold, UtrCH–CF555, and H2A–H2B chromobody–mScarlett3 to label microtubules (white), actin (cyan), and DNA (magenta), respectively, under control conditions (C) or following treatment with 20 µM nocodazole (D). The yellow arrowhead indicates the fertilization site, and white arrows mark the aster center (male pronucleus). Scale bar, 10 µm. (E) Dynamics of the average distance between the aster center and the cell cortex (left axis) and the average maximal actin cable length at the fertilization site (right axis) in control zygotes (n = 6). Dotted lines indicate the standard error.

**Figure S4:**
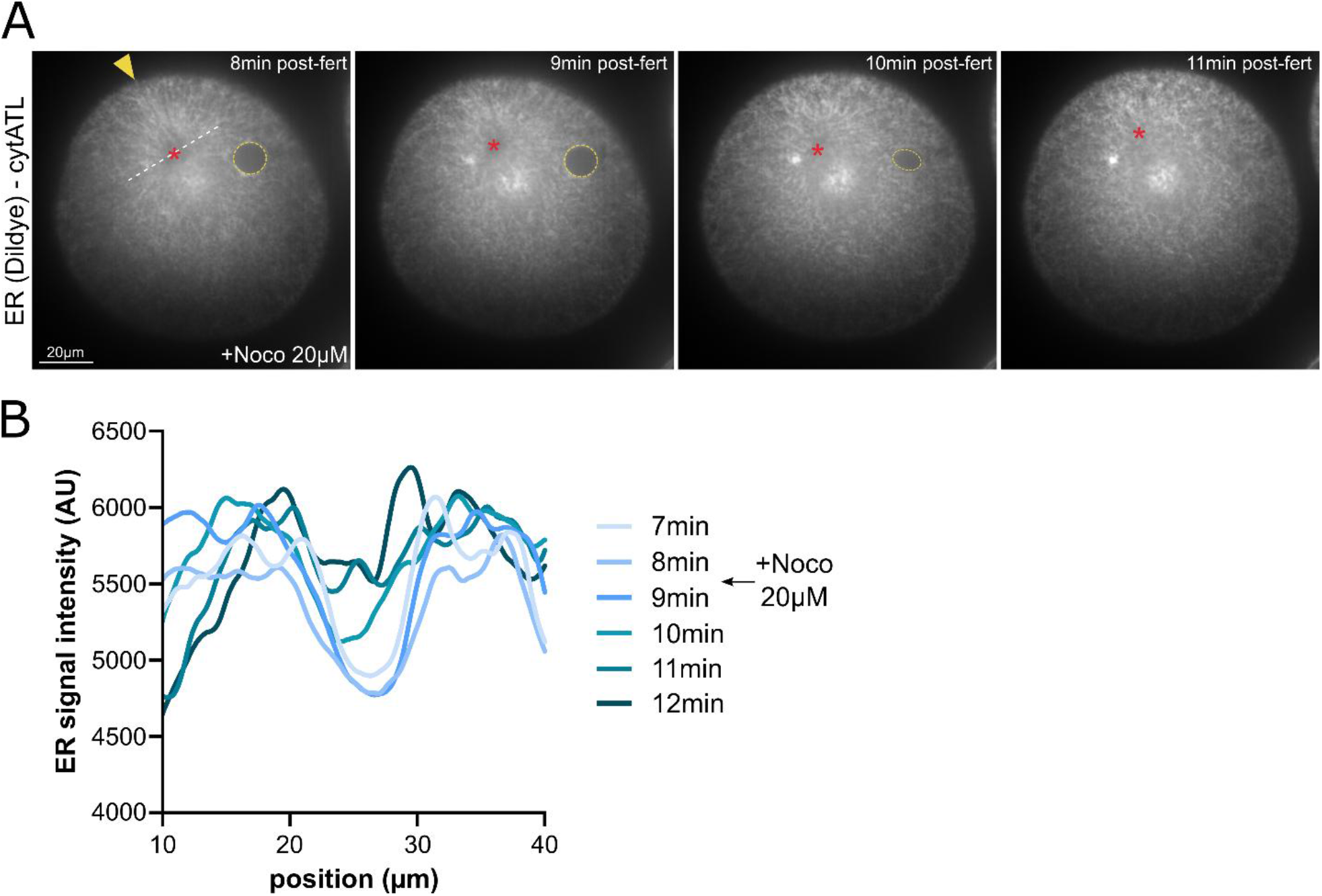
ER fragmentation in cytATL injected zygotes is microtubule dependent. (A) Time-lapse images of aster centration following DiIdye injection (ER labeling) and cytATL injection, in a zygote treated with 20 µM nocodazole 8 min after fertilization. The yellow arrowhead indicates the fertilization site, the red star marks the aster center, the yellow dotted circle outlines the female pronucleus, and the white dotted line indicates the region where the intensity profile was measured. Scale bar, 20 µm. (B) Corresponding ER intensity profiles over time.

## MOVIE LEGENDS

**Movie1:** Single-plane time-lapse imaging of sperm aster centration acquired using an Airyscan microscope. Microtubules are shown in white (injected TMBD–mStayGold protein), and actin is shown in cyan (injected UtrCH–CF555 protein). Frames were acquired every 2 s. The signal-to-noise ratio was enhanced using a 3-frame temporal averaging.

**Movie2:** Single plane time-lapse imaging of sperm aster centration acquired using an Airyscan confocal microscope. Microtubules are in white (injected TMBD-mStayGold protein). Yellow arrows are showing microtubules detaching from the aster. Frames were acquired every second. The signal-to-noise ratio was enhanced using a three-frame temporal averaging.

**Movie3:** Time-lapse imaging of the “stalled aster” assay captured on a widefield microscope (DIC). Yellow arrow shows the position of the magnetic particles aggregate. Frames were acquired every second.

**Movie4:** Independent single-plane time-lapse imaging of lysosomes (LysoTracker), endosomes (FM1-43), mitochondria (MitoTracker) and yolk granules (Nile blue) during aster centration acquired on a spinning-disc confocal microscope. The red star indicates the location of the aster center. Frames were acquired every 2 s.

**Movie5:** Single plane time-lapse imaging of endoplasmic reticulum (DilDye injection) during sperm aster centration acquired using a spinning-disc confocal microscope. Frames were acquired every minute.

**Movie6:** Single-plane, high–frame rate time-lapse imaging of the endoplasmic reticulum (DilDye injection) during sperm aster centration, acquired using a spinning-disc confocal microscope. Frames were acquired every 2 s.

**Movie7**: Successive time-lapse imaging of pronuclei motion (Hoechst 33342) during aster centration after cytATL or cytATL^R232Q^ protein injection acquired on a wide-field fluorescence microscope. Frames were acquired every minute.

**Movie8**: Single plane time-lapse imaging of sperm aster centration after cytATL protein injection acquired using a spinning-disc confocal microscope. Microtubules were labelled using TMBD-mStayGold protein. Frames were acquired every minute.

**Movie9**: Single plane time-lapse imaging of endoplasmic reticulum (DilDye injection) during sperm aster centration after cytATL protein injection acquired using a spinning-disc confocal microscope. Frames were acquired every minute.

**Movie10**: Single-plane, high–frame rate time-lapse imaging of the endoplasmic reticulum (DilDye injection) during sperm aster centration after cytATL protein injection acquired using a spinning-disc confocal microscope. Yellow arrows show examples of membrane tubes being pulled from the endoplasmic reticulum and the nuclear envelope. Frames were acquired every second.

**Movie11**: Single-plane, high–frame rate time-lapse imaging of the endoplasmic reticulum (DilDye injection) during 1 cell stage metaphase after cytATL protein injection acquired using a spinning-disc confocal microscope. Yellow arrows show examples of membrane tubes being pulled from the endoplasmic reticulum. Red stars show the location of spindle poles. Frames were acquired every second.

**MovieS1**: Single-plane, high–frame rate time-lapse imaging of microtubule plus ends (EB1-GFP protein injection) during aster centration acquired using a spinning-disc confocal microscope. Frames were acquired every second. The signal-to-noise ratio was enhanced using a two-frame temporal averaging.

**MovieS2**: Time-lapse imaging of pronuclear movements during aster centration in (i) a non-injected zygote, (ii) a buffer-injected zygote, (iii) a zygote injected with p150^CC1^ exhibiting failed centration, (iv) a zygote injected with p150^CC1^ exhibiting overshoot centration, (v) a zygote injected with *P.liv* p150^CC1^ exhibiting failed centration, or (vi) a zygote injected with *P.liv* p150^CC1^ exhibiting overshoot centration. Images were acquired using a wide-field fluorescence microscope. The DIC channel is shown in white, and DNA (Hoechst 33342) is shown in cyan. Frames were acquired every minute.

**MovieS3**: Time-lapse imaging of pronuclei motion during early aster centration in (i) a control condition (DMSO), (ii) a zygote treated with 20µM Nocodazole, (iii) a zygote treated with 20µM Latrunculin B, or (iv) a zygote treated with both 20µM Nocodazole and 20µM Latrunculin B. Images were acquired on a wide-field fluorescence microscope. DIC channel is white and DNA (Hoechst 33342) is in cyan. Frames were acquired every 30 s.

**MovieS4**: Time-lapse imaging of pronuclei motion during early aster centration in a control condition (DMSO), acquired on a spinning-disc confocal microscope. Microtubules are in orange (injected TMBD–mStayGold protein), actin is in cyan (injected UtrCH–CF555 protein) and DNA is in magenta (injected H2A-H2B-chromobody-mScarlett3). Frames were acquired every 30 s.

**MovieS5**: Time-lapse imaging of pronuclei motion during early aster centration in a zygote treated with 20µM Nocodazole, acquired on a spinning-disc confocal microscope. Microtubules are in orange (injected TMBD–mStayGold protein), actin in cyan (injected UtrCH–CF555 protein) and DNA in magenta (injected H2A-H2B-chromobody-mScarlett3). Frames were acquired every 30 s.

**MovieS6**: Single plane time-lapse imaging of endoplasmic reticulum (DilDye injection) during sperm aster centration, acquired using a spinning-disc confocal microscope. The label “+Noco” indicates the frames at which 20 µM nocodazole was added. Frames were acquired every minute.

## REFERENCES

1. Sallé J, Minc N. Cell division geometries as central organizers of early embryo development. Seminars in Cell & Developmental Biology. 2022 Oct 1;Special issue: The control of tissue size and shape by Yanlan Mao and Franck Pichauds 130:3–11. doi:10.1016/j.semcdb.2021.08.004

2. Minc N, Burgess D, Chang F. Influence of cell geometry on division-plane positioning. Cell. 2011 Feb 4;144(3):414–26. doi:10.1016/j.cell.2011.01.016 PubMed PMID: 21295701; PubMed Central PMCID: PMC3048034.

3. Wu HY, Nazockdast E, Shelley MJ, Needleman DJ. Forces positioning the mitotic spindle: Theories, and now experiments. Bioessays. 2017;39(2). doi:10.1002/bies.201600212 PubMed PMID: 28026040.

4. Kroll J, Renkawitz J. Principles of organelle positioning in motile and non-motile cells. EMBO reports. 2024 May 14;25(5):2172–87. doi:10.1038/s44319-024-00135-4

5. Reinsch S, Gonczy P. Mechanisms of nuclear positioning. J Cell Sci. 1998 Aug;111 (Pt 16):2283–95.

6. Tran PT, Marsh L, Doye V, Inoué S, Chang F. A Mechanism for Nuclear Positioning in Fission Yeast Based on Microtubule Pushing. The Journal of Cell Biology. 2001 Apr 16;153(2):397–412. doi:10.1083/jcb.153.2.397 PubMed PMID: 11309419.

7. Dogterom M, Kerssemakers JW, Romet-Lemonne G, Janson ME. Force generation by dynamic microtubules. Curr Opin Cell Biol. 2005 Feb;17(1):67–74. doi:10.1016/j.ceb.2004.12.011

8. Meaders JL, de Matos SN, Burgess DR. A Pushing Mechanism for Microtubule Aster Positioning in a Large Cell Type. Cell Rep. 2020 Oct 6;33(1):108213. doi:10.1016/j.celrep.2020.108213 PubMed PMID: 33027648.

9. Garzon-Coral C, Fantana HA, Howard J. A force-generating machinery maintains the spindle at the cell center during mitosis. Science. 2016 May 27;352(6289):1124–7. doi:10.1126/science.aad9745 PubMed PMID: 27230381.

10. Laan L, Pavin N, Husson J, Romet-Lemonne G, van Duijn M, López MP, et al. Cortical dynein controls microtubule dynamics to generate pulling forces that position microtubule asters. Cell. 2012 Feb 3;148(3):502–14. doi:10.1016/j.cell.2012.01.007 PubMed PMID: 22304918; PubMed Central PMCID: PMC3292199.

11. Grill SW, Gönczy P, Stelzer EH, Hyman AA. Polarity controls forces governing asymmetric spindle positioning in the Caenorhabditis elegans embryo. Nature. 2001 Feb 1;409(6820):630–3. doi:10.1038/35054572 PubMed PMID: 11214323.

12. Labbé JC, McCarthy EK, Goldstein B. The forces that position a mitotic spindle asymmetrically are tethered until after the time of spindle assembly. The Journal of Cell Biology. 2004 Oct 25;167(2):245–56. doi:10.1083/jcb.200406008 PubMed PMID: 15492042.

13. Hernández-López C, Puliafito A, Xu Y, Lu Z, Di Talia S, Vergassola M. Two-fluid dynamics and micron-thin boundary layers shape cytoplasmic flows in early Drosophila embryos. Proc Natl Acad Sci U S A. 2023 Oct 31;120(44):e2302879120. doi:10.1073/pnas.2302879120 PubMed PMID: 37878715; PubMed Central PMCID: PMC10622894.

14. Shamipour S, Kardos R, Xue SL, Hof B, Hannezo E, Heisenberg CP. Bulk Actin Dynamics Drive Phase Segregation in Zebrafish Oocytes. Cell. 2019 May 30;177(6):1463–1479.e18. doi:10.1016/j.cell.2019.04.030

15. Xie J, Minc N. Cytoskeleton Force Exertion in Bulk Cytoplasm. Front Cell Dev Biol. 2020 Feb 13;8. doi:10.3389/fcell.2020.00069

16. Hamaguchi MS, Hiramoto Y. Analysis of the Role of Astral Rays in Pronuclear Migration in Sand Dollar Eggs by the Colcemid-UV Method. Development, Growth & Differentiation. 1986 Apr 1;28(2):143–56. doi:10.1111/j.1440-169X.1986.00143.x

17. Caballero-Mancebo S, Suarez DG, Chenevert J, Ben-Aicha S, Besnardeau L, McDougall A, et al. Cooperation between cortical and cytoplasmic forces shapes planar 4-cell stage embryos [Internet]. bioRxiv; 2025 [cited 2026 Jun 1]. p. 2025.11.24.690087. Available from: https://www.biorxiv.org/content/10.1101/2025.11.24.690087v3 doi:10.1101/2025.11.24.690087

18. Kiyomitsu A, Bai L, Mitchison TJ, Kiyomitsu T. Spatiotemporal organization of dynein in bulk cytoplasm promotes aster growth and positioning in large embryos [Internet]. bioRxiv; 2026 [cited 2026 Apr 7]. p. 2026.04.02.716240. Available from: https://www.biorxiv.org/content/10.64898/2026.04.02.716240v1 doi:10.64898/2026.04.02.716240

19. Pelletier JF, Field CM, Fürthauer S, Sonnett M, Mitchison TJ. Co-movement of astral microtubules, organelles and F-actin by dynein and actomyosin forces in frog egg cytoplasm. Surrey T, Malhotra V, Di Talia S, editors. eLife. 2020 Dec 7;9:e60047. doi:10.7554/eLife.60047

20. Cheng X, Ferrell JE. Spontaneous emergence of cell-like organization in Xenopus egg extracts. Science. 2019 01;366(6465):631–7. doi:10.1126/science.aav7793 PubMed PMID: 31672897.

21. Rinaldin M, Kickuth A, Lamson A, Dalton B, Xu Y, Mejstřík P, et al. Robust cytoplasmic partitioning by solving a cytoskeletal instability. Nature. 2026 Mar;651(8105):501–10. doi:10.1038/s41586-025-10023-z

22. Bai L, Field CM, Kiyomitsu A, Shen Y, Orlovsky ND, Kiyomitsu T, et al. Local fluidization of an active cytoplasmic gel partitions large cells [Internet]. bioRxiv; 2026 [cited 2026 Sep 8]. p. 2026.08.11.744254. Available from: https://www.biorxiv.org/content/10.64898/2026.08.11.744254v1 doi:10.64898/2026.08.11.744254

23. Kimura K, Kimura A. Intracellular organelles mediate cytoplasmic pulling force for centrosome centration in the Caenorhabditis elegans early embryo. Proc Natl Acad Sci USA. 2011 Jan 4;108(1):137–42. doi:10.1073/pnas.1013275108 PubMed PMID: 21173218; PubMed Central PMCID: PMC3017145.

24. Hamaguchi MS, Hamaguchi Y, Hiramoto Y. Microinjected Polystyrene Beads Move Along Astral Rays in Sand Dollar Eggs. Development, Growth & Differentiation. 1986;28(5):461–70. doi:10.1111/j.1440-169X.1986.00461.x

25. Tanimoto H, Sallé J, Dodin L, Minc N. Physical Forces Determining the Persistency and Centering Precision of Microtubule Asters. Nat Phys. 2018 Aug;14(8):848–54. doi:10.1038/s41567-018-0154-4 PubMed PMID: 30079097; PubMed Central PMCID: PMC6071857.

26. Tanimoto H, Kimura A, Minc N. Shape-motion relationships of centering microtubule asters. J Cell Biol. 2016 Mar 28;212(7):777–87. doi:10.1083/jcb.201510064 PubMed PMID: 27022090; PubMed Central PMCID: PMC4810306.

27. Kimura A, Onami S. Computer Simulations and Image Processing Reveal Length-Dependent Pulling Force as the Primary Mechanism for C. elegans Male Pronuclear Migration. Developmental Cell. 2005 May 1;8(5):765–75. doi:10.1016/j.devcel.2005.03.007 PubMed PMID: 15866166.

28. Wuhr M, Tan ES, Parker SK, Detrich HW, Mitchison TJ. A model for cleavage plane determination in early amphibian and fish embryos. Curr Biol. 2010 Nov 23;20(22):2040–5. doi:10.1016/j.cub.2010.10.024

29. Rosfelter A, de Labbey G, Chenevert J, Dumollard R, Schaub S, Machaty Z, et al. Reduction of cortical pulling at mitotic entry facilitates aster centration. Journal of Cell Science. 2024 Mar 12;jcs.262037. doi:10.1242/jcs.262037

30. De Simone A, Spahr A, Busso C, Gönczy P. Uncovering the balance of forces driving microtubule aster migration in C. elegans zygotes. Nature Communications. 2018 Mar 5;9(1):938. doi:10.1038/s41467-018-03118-x

31. Barbosa DJ, Duro J, Prevo B, Cheerambathur DK, Carvalho AX, Gassmann R. Dynactin binding to tyrosinated microtubules promotes centrosome centration in C. elegans by enhancing dynein-mediated organelle transport. PLOS Genetics. 2017 Jul 31;13(7):e1006941. doi:10.1371/journal.pgen.1006941

32. Afonso O, Dumoulin L, Kruse K, Gonzalez-Gaitan M. Cytoplasmic flow is a cell size sensor that scales anaphase. Nat Cell Biol. 2025 Jan 31;1–10. doi:10.1038/s41556-024-01605-6

33. Shinar T, Mana M, Piano F, Shelley MJ. A model of cytoplasmically driven microtubule-based motion in the single-celled Caenorhabditis elegans embryo. Proc Natl Acad Sci USA. 2011 Jun 28;108(26):10508–13. doi:10.1073/pnas.1017369108 PubMed PMID: 21670261; PubMed Central PMCID: PMC3127902.

34. Longoria RA, Shubeita GT. Cargo Transport by Cytoplasmic Dynein Can Center Embryonic Centrosomes. PLOS ONE. 2013 Jul 1;8(7):e67710. doi:10.1371/journal.pone.0067710

35. Letort G, Nedelec F, Blanchoin L, Th éry M, Mogilner A. Centrosome centering and decentering by microtubule network rearrangement. Molecular Biology of the Cell. 2016 Sep 15;27(18):2833–43. doi:10.1091/mbc.E16-06-0395

36. Palenzuela H, Lacroix B, Sallé J, Minami K, Shima T, Jegou A, et al. In vitro reconstitution of dynein force exertion in a bulk viscous medium. Current Biology. 2020;30(22):4534–4540. e7.

37. Nazockdast E, Rahimian A, Needleman D, Shelley M. Cytoplasmic flows as signatures for the mechanics of mitotic positioning. Mol Biol Cell. 2017 Nov 7;28(23):3261–70. doi:10.1091/mbc.E16-02-0108

38. Sulerud T, Sami AB, Li G, Kloxin A, Oakey J, Gatlin J. Microtubule-dependent pushing forces contribute to long-distance aster movement and centration in Xenopus laevis egg extracts. Mol Biol Cell. 2020 Dec 1;31(25):2791–802. doi:10.1091/mbc.E20-01-0088 PubMed PMID: 33026931; PubMed Central PMCID: PMC7851858.

39. Chambers EL. The Movement of the Egg Nucleus in Relation to the Sperm Aster in the Echinoderm Egg. Journal of Experimental Biology. 1939;16(4):409–24.

40. Schatten H, Schatten G, Mazia D, Balczon R, Simerly C. Behavior of centrosomes during fertilization and cell division in mouse oocytes and in sea urchin eggs. Proc Natl Acad Sci U S A. 1986 Jan;83(1):105–9.

41. Mooney P, Sulerud T, Pelletier JF, Dilsaver MR, Tomschik M, Geisler C, et al. Tau-based fluorescent protein fusions to visualize microtubules. Cytoskeleton (Hoboken). 2017 Jun;74(6):221–32. doi:10.1002/cm.21368 PubMed PMID: 28407416; PubMed Central PMCID: PMC5592782.

42. Ivorra-Molla E, Akhuli D, McAndrew MBL, Scott W, Kumar L, Palani S, et al. A monomeric StayGold fluorescent protein. Nat Biotechnol. 2023 Dec 11;1–4. doi:10.1038/s41587-023-02018-w

43. Tirado MM, de la Torre JG. Translational friction coefficients of rigid, symmetric top macromolecules. Application to circular cylinders. J Chem Phys. 1979 Sep 15;71(6):2581–7. doi:10.1063/1.438613

44. Quintyne NJ, Gill SR, Eckley DM, Crego CL, Compton DA, Schroer TA. Dynactin is required for microtubule anchoring at centrosomes. J Cell Biol. 1999 Oct 18;147(2):321–34. doi:10.1083/jcb.147.2.321 PubMed PMID: 10525538; PubMed Central PMCID: PMC2174233.

45. Reinsch S, Karsenti E. Movement of nuclei along microtubules in Xenopus egg extracts. Current Biology. 1997 Mar 1;7(3):211–4. doi:10.1016/S0960-9822(97)70092-7

46. Chun JT, Vasilev F, Limatola N, Santella L. Fertilization in Starfish and Sea Urchin: Roles of Actin. In: Kloc M, Kubiak JZ, editors. Marine Organisms as Model Systems in Biology and Medicine [Internet]. Cham: Springer International Publishing; 2018 [cited 2025 Sep 19]. p. 33–47. Available from: 10.1007/978-3-319-92486-1_3 doi:10.1007/978-3-319-92486-1_3

47. Scheffler K, Uraji J, Jentoft I, Cavazza T, Mönnich E, Mogessie B, et al. Two mechanisms drive pronuclear migration in mouse zygotes. Nat Commun. 2021 Feb 5;12(1):841. doi:10.1038/s41467-021-21020-x

48. Jimenez AJ, Schaeffer A, De Pascalis C, Letort G, Vianay B, Bornens M, et al. Acto-myosin network geometry defines centrosome position. Current Biology. 2021 Mar 22;31(6):1206–1220.e5. doi:10.1016/j.cub.2021.01.002

49. Hirohashi N, Yanagimachi R. Sperm acrosome reaction: its site and role in fertilization†. Biol Reprod. 2018 Jul 1;99(1):127–33. doi:10.1093/biolre/ioy045

50. Terasaki M. Actin filament translocations in sea urchin eggs. Cell Motil Cytoskeleton. 1996;34(1):48–56. doi:10.1002/(SICI)1097-0169(1996)34:1<48::AID-CM5>3.0.CO;2-E PubMed PMID: 8860231.

51. Xie J, Najafi J, Le Borgne R, Verbavatz JM, Durieu C, Sallé J, et al. Contribution of cytoplasm viscoelastic properties to mitotic spindle positioning. Proceedings of the National Academy of Sciences. 2022 Feb 22;119(8):e2115593119. doi:10.1073/pnas.2115593119

52. Najafi J, Dmitrieff S, Minc N. Size- and position-dependent cytoplasm viscoelasticity through hydrodynamic interactions with the cell surface. Proceedings of the National Academy of Sciences. 2023 Feb 28;120(9):e2216839120. doi:10.1073/pnas.2216839120

53. Sallé J, Xie J, Ershov D, Lacassin M, Dmitrieff S, Minc N. Asymmetric division through a reduction of microtubule centering forces. J Cell Biol. 2019 Mar 4;218(3):771–82. doi:10.1083/jcb.201807102 PubMed PMID: 30563876; PubMed Central PMCID: PMC6400563.

54. English AR, Zurek N, Voeltz GK. Peripheral ER structure and function. Curr Opin Cell Biol. 2009 Aug;21(4):596–602. doi:10.1016/j.ceb.2009.04.004 PubMed PMID: 19447593; PubMed Central PMCID: PMC2753178.

55. Arjona MI, Khosravanizadeh A, Municio-Díaz C, Mioche M, Dmitrieff S, Sallé J, et al. A Composite Endomembrane Suspension Governs Cytoplasm Rheology [Internet]. bioRxiv; 2026 [cited 2026 Sep 8]. p. 2026.07.20.739612. Available from: https://www.biorxiv.org/content/10.64898/2026.07.20.739612v1 doi:10.64898/2026.07.20.739612

56. Kimura K, Mamane A, Sasaki T, Sato K, Takagi J, Niwayama R, et al. Endoplasmic-reticulum-mediated microtubule alignment governs cytoplasmic streaming. Nat Cell Biol. 2017 Apr;19(4):399–406. doi:10.1038/ncb3490

57. Terasaki M, Jaffe LA. Organization of the sea urchin egg endoplasmic reticulum and its reorganization at fertilization. J Cell Biol. 1991 Sep;114(5):929–40. PubMed PMID: 1874789; PubMed Central PMCID: PMC2289104.

58. Mukherjee RN, Sallé J, Dmitrieff S, Nelson KM, Oakey J, Minc N, et al. The Perinuclear ER Scales Nuclear Size Independently of Cell Size in Early Embryos. Dev Cell. 2020 Aug 10;54(3):395–409.e7. doi:10.1016/j.devcel.2020.05.003 PubMed PMID: 32473090; PubMed Central PMCID: PMC7423768.

59. Rai AK, Rai A, Ramaiya AJ, Jha R, Mallik R. Molecular Adaptations Allow Dynein to Generate Large Collective Forces inside Cells. Cell. 2013 Jan 17;152(1):172–82. doi:10.1016/j.cell.2012.11.044 PubMed PMID: 23332753.

60. Immers J. Studies on cytoplasmic components of sea urchin eggs stratified by centrifugation. Experimental Cell Research. 1960 Apr 1;19(3):499–514. doi:10.1016/0014-4827(60)90059-8

61. Gross PR, Philpott DE, Nass S. Electron microscopy of the centrifuged sea urchin egg, with a note on the structure of the ground cytoplasm. J Biophys Biochem Cytol. 1960 Feb;7(1):135–42. doi:10.1083/jcb.7.1.135 PubMed PMID: 13829660; PubMed Central PMCID: PMC2224862.

62. Wang N, Rapoport TA. Reconstituting the reticular ER network – mechanistic implications and open questions. Journal of Cell Science. 2019 Jan 22;132(4):jcs227611. doi:10.1242/jcs.227611

63. Wang S, Romano FB, Field CM, Mitchison TJ, Rapoport TA. Multiple mechanisms determine ER network morphology during the cell cycle in Xenopus egg extracts. J Cell Biol. 2013 Dec 2. doi:10.1083/jcb.201308001

64. Wang S, Tukachinsky H, Romano FB, Rapoport TA. Cooperation of the ER-shaping proteins atlastin, lunapark, and reticulons to generate a tubular membrane network. Elife. 2016 Sep 13;5. doi:10.7554/eLife.18605

65. Farías GG, Fréal A, Tortosa E, Stucchi R, Pan X, Portegies S, et al. Feedback-Driven Mechanisms between Microtubules and the Endoplasmic Reticulum Instruct Neuronal Polarity. Neuron. 2019 Apr 3;102(1):184–201.e8. doi:10.1016/j.neuron.2019.01.030 PubMed PMID: 30772082.

66. Terasaki M, Chen LB, Fujiwara K. Microtubules and the endoplasmic reticulum are highly interdependent structures. J Cell Biol. 1986 Oct;103(4):1557–68. doi:10.1083/jcb.103.4.1557 PubMed PMID: 3533956; PubMed Central PMCID: PMC2114338.

67. Waterman-Storer CM, Salmon ED. Endoplasmic reticulum membrane tubules are distributed by microtubules in living cells using three distinct mechanisms. Curr Biol. 1998 Jul 2;8(14):798–806. doi:10.1016/s0960-9822(98)70321-5 PubMed PMID: 9663388.

68. Leduc C, Campàs O, Zeldovich KB, Roux A, Jolimaitre P, Bourel-Bonnet L, et al. Cooperative extraction of membrane nanotubes by molecular motors. Proceedings of the National Academy of Sciences. 2004 Dec 7;101(49):17096–101. doi:10.1073/pnas.0406598101

69. Scott ZC, Steen SB, Huber G, Westrate LM, Koslover EF. The endoplasmic reticulum as an active liquid network. Proceedings of the National Academy of Sciences. 2024 Oct 15;121(42):e2409755121. doi:10.1073/pnas.2409755121

70. Teixeira V, Singh K, Gama JB, Moreira M, Celestino R, Xavier Carvalho A, et al. CDR2 is a dynein adaptor recruited by kinectin to regulate ER sheet organization. J Cell Biol. 2025 Jul 10;224(9):e202411034. doi:10.1083/jcb.202411034

71. Allan V. Protein phosphatase 1 regulates the cytoplasmic dynein-driven formation of endoplasmic reticulum networks in vitro. J Cell Biol. 1995 Mar 1;128(5):879–91. doi:10.1083/jcb.128.5.879

72. Steffen W, Karki S, Vaughan KT, Vallee RB, Holzbaur ELF, Weiss DG, et al. The Involvement of the Intermediate Chain of Cytoplasmic Dynein in Binding the Motor Complex to Membranous Organelles ofXenopus Oocytes. Molecular Biology of the Cell. 1997 Oct;8(10):2077–88. doi:10.1091/mbc.8.10.2077

73. Kimura K, Kimura A. A novel mechanism of microtubule length-dependent force to pull centrosomes toward the cell center. BioArchitecture. 2014;1(2):74–9. doi:10.4161/bioa.1.2.15549

74. Alnæs M, Blechta J, Hake J, Johansson A, Kehlet B, Logg A, et al. The FEniCS Project Version 1.5. Archive of Numerical Software. 2015 Dec 7;3(100). doi:10.11588/ans.2015.100.20553

75. Logg A, Mardal KA, Wells G, editors. Automated Solution of Differential Equations by the Finite Element Method: The FEniCS Book [Internet]. Vol. 84. Berlin, Heidelberg: Springer; 2012 [cited 2026 May 27]. (Lecture Notes in Computational Science and Engineering). Available from: https://link.springer.com/10.1007/978-3-642-23099-8 doi:10.1007/978-3-642-23099-8

76. Pierre A, Salle J, Wuhr M, Minc N. Generic Theoretical Models to Predict Division Patterns of Cleaving Embryos. Dev Cell. 2016 Dec 19;39(6):667–82. doi:10.1016/j.devcel.2016.11.018

77. Godard BG, Dumollard R, Heisenberg CP, McDougall A. Combined effect of cell geometry and polarity domains determines the orientation of unequal division. Devenport D, Akhmanova A, editors. eLife. 2021 Dec 10;10:e75639. doi:10.7554/eLife.75639

78. Niwayama R, Moghe P, Liu YJ, Fabrèges D, Buchholz F, Piel M, et al. A Tug-of-War between Cell Shape and Polarity Controls Division Orientation to Ensure Robust Patterning in the Mouse Blastocyst. Developmental Cell. 2019 Dec 2;51(5):564–574.e6. doi:10.1016/j.devcel.2019.10.012

79. Suzuki K, Miyazaki M, Takagi J, Itabashi T, Ishiwata S. Spatial confinement of active microtubule networks induces large-scale rotational cytoplasmic flow. Proceedings of the National Academy of Sciences. 2017 Mar 14;114(11):2922–7. Located at: world. doi:10.1073/pnas.1616001114

80. Lane JD, Allan VJ. Microtubule-based endoplasmic reticulum motility in Xenopus laevis: activation of membrane-associated kinesin during development. Mol Biol Cell. 1999 Jun;10(6):1909–22. doi:10.1091/mbc.10.6.1909 PubMed PMID: 10359605; PubMed Central PMCID: PMC25389.

81. Woźniak MJ, Bola B, Brownhill K, Yang YC, Levakova V, Allan VJ. Role of kinesin-1 and cytoplasmic dynein in endoplasmic reticulum movement in VERO cells. J Cell Sci. 2009 Jun 15;122(12):1979–89. doi:10.1242/jcs.041962 PubMed PMID: 19454478; PubMed Central PMCID: PMC2723153.

82. Karabasheva D, Smyth JT. A novel, dynein-independent mechanism focuses the endoplasmic reticulum around spindle poles in dividing Drosophila spermatocytes. Scientific Reports. 2019 Aug 28;9(1):1. doi:10.1038/s41598-019-48860-4

83. Watson P, Forster R, Palmer KJ, Pepperkok R, Stephens DJ. Coupling of ER exit to microtubules through direct interaction of COPII with dynactin. Nat Cell Biol. 2005 Jan;7(1):48–55. doi:10.1038/ncb1206

84. Zheng P, Obara CJ, Szczesna E, Nixon-Abell J, Mahalingan KK, Roll-Mecak A, et al. ER proteins decipher the tubulin code to regulate organelle distribution. Nature. 2022 Jan;601(7891):132–8. doi:10.1038/s41586-021-04204-9 PubMed PMID: 34912111; PubMed Central PMCID: PMC8732269.

85. Grigoriev I, Gouveia SM, van der Vaart B, Demmers J, Smyth JT, Honnappa S, et al. STIM1 is a MT-plus-end-tracking protein involved in remodeling of the ER. Curr Biol. 2008 Feb 12;18(3):177–82. doi:10.1016/j.cub.2007.12.050 PubMed PMID: 18249114; PubMed Central PMCID: PMC2600655.

86. Harvey EB. Effects of centrifugal force on the ectoplasmic layer and nuclei of fertilized sea urchin eggs. The Biological Bulletin. 1934 Apr;66(2):228–45. doi:10.2307/1537334

87. Anderson E. A Cytological Study of the Centrifuged Whole, Half, and Quarter Eggs of the Sea Urchin Arbacia Punctulata. The Journal of Cell Biology. 1970 Dec 1;47(3):711–33. doi:10.1083/jcb.47.3.711 PubMed PMID: 5497548.

88. Henson JH, Begg DA, Beaulieu SM, Fishkind DJ, Bonder EM, Terasaki M, et al. A calsequestrin-like protein in the endoplasmic reticulum of the sea urchin: localization and dynamics in the egg and first cell cycle embryo. J Cell Biol. 1989 Jul;109(1):149–61. doi:10.1083/jcb.109.1.149 PubMed PMID: 2663877; PubMed Central PMCID: PMC2115469.

89. Xie J, Levy DL, Minc N, Sallé J. Manipulation of Embryonic Cleavage Geometry Using Magnetic Tweezers. Methods Mol Biol. 2024;2740:125–40. doi:10.1007/978-1-0716-3557-5_8 PubMed PMID: 38393473.

90. Ershov D, Phan MS, Pylvänäinen JW, Rigaud SU, Le Blanc L, Charles-Orszag A, et al. TrackMate 7: integrating state-of-the-art segmentation algorithms into tracking pipelines. Nat Methods. 2022 Jul;19(7):829–32. doi:10.1038/s41592-022-01507-1

91. Berg S, Kutra D, Kroeger T, Straehle CN, Kausler BX, Haubold C, et al. ilastik: interactive machine learning for (bio)image analysis. Nat Methods. 2019 Dec;16(12):1226–32. doi:10.1038/s41592-019-0582-9 PubMed PMID: 31570887.

92. Gros O, Bhickta C, Lokaj G, Schwab Y, Köhler S, Banterle N. Microscopy Nodes: versatile 3D microscopy visualization with Blender [Internet]. bioRxiv; 2025 [cited 2025 Apr 29]. p. 2025.01.09.632153. Available from: https://www.biorxiv.org/content/10.1101/2025.01.09.632153v1 doi:10.1101/2025.01.09.632153

93. Bun P, Dmitrieff S, Belmonte JM, Nédélec FJ, Lénárt P. A disassembly-driven mechanism explains F-actin-mediated chromosome transport in starfish oocytes. eLife. 2018;7:e31469. doi:10.7554/eLife.31469 PubMed PMID: 29350616; PubMed Central PMCID: PMC5788506.

94. Schlager MA, Serra-Marques A, Grigoriev I, Gumy LF, Esteves da Silva M, Wulf PS, et al. Bicaudal d family adaptor proteins control the velocity of Dynein-based movements. Cell Rep. 2014 Sep 11;8(5):1248–56. doi:10.1016/j.celrep.2014.07.052 PubMed PMID: 25176647.

95. McKenney RJ, Huynh W, Tanenbaum ME, Bhabha G, Vale RD. Activation of cytoplasmic dynein motility by dynactin-cargo adapter complexes. Science. 2014 Jul 18;345(6194):337–41. doi:10.1126/science.1254198 PubMed PMID: 25035494; PubMed Central PMCID: PMC4224444.

96. Schroeder CM, Vale RD. Assembly and activation of dynein-dynactin by the cargo adaptor protein Hook3. J Cell Biol. 2016 Aug 1;214(3):309–18. doi:10.1083/jcb.201604002 PubMed PMID: 27482052; PubMed Central PMCID: PMC4970328.

97. Howard J. Mechanics of Motor Proteins and the Cytoskeleton. Sinauer Associates, Publishers; 2001. 367 p.

98. Geuzaine C, Remacle JF. Gmsh: A 3-D finite element mesh generator with built-in pre- and post-processing facilities. International Journal for Numerical Methods in Engineering. 2009;79(11):1309–31. doi:10.1002/nme.2579

99. Toba S, Watanabe TM, Yamaguchi-Okimoto L, Toyoshima YY, Higuchi H. Overlapping hand-over-hand mechanism of single molecular motility of cytoplasmic dynein. Proceedings of the National Academy of Sciences. 2006 Apr 11;103(15):5741–5. doi:10.1073/pnas.0508511103

